# Design and power analysis for multi-sample single cell genomics experiments

**DOI:** 10.1101/2020.04.01.019851

**Authors:** Katharina T. Schmid, Cristiana Cruceanu, Anika Böttcher, Heiko Lickert, Elisabeth B. Binder, Fabian J. Theis, Matthias Heinig

## Abstract

**Background:** The identification of genes associated with specific experimental conditions, genotypes or phenotypes through differential expression analysis has long been the cornerstone of transcriptomic analysis. Single cell RNA-seq is revolutionizing transcriptomics and is enabling interindividual differential gene expression analysis and identification of genetic variants associated with gene expression, so called expression quantitative trait loci at cell-type resolution. Current methods for power analysis and guidance of experimental design either do not account for the specific characteristics of single cell data or are not suitable to model interindividual comparisons.

**Results:** Here we present a statistical framework for experimental design and power analysis of single cell differential gene expression between groups of individuals and expression quantitative trait locus analysis. The model relates sample size, number of cells per individual and sequencing depth to the power of detecting differentially expressed genes within individual cell types. Power analysis is based on data driven priors from literature or pilot experiments across a wide range of application scenarios and single cell RNA-seq platforms. Using these priors we show that, for a fixed budget, the number of cells per individual is the major determinant of power.

**Conclusion:** Our model is general and allows for systematic comparison of alternative experimental designs and can thus be used to guide experimental design to optimize power. For a wide range of applications, shallow sequencing of high numbers of cells per individual leads to higher overall power than deep sequencing of fewer cells. The model is implemented as an R package *scPower*.

## Background

From the early days of microarrays, one of the main goals of transcriptomic profiling has been to identify changes of gene expression levels (differentially expressed genes; DEGs) between sets of samples, e.g. patients and healthy controls [1-5]. With the advent of single cell RNA-sequencing (scRNA-seq) [6-10], the sets of samples denote sets of cells of a particular type. In the context of single cell genomics, cell types or states describe the cellular phenotype in terms of its expression profile and are typically derived from the data [11]. Here we focus on a discrete notion of cell types that is typically derived by clustering [12], potentially on different levels of granularity [13]. Single cell differential gene expression analysis typically seeks to identify genes whose expression levels are markedly different between sets of cells of different cell types [14-16]. Here, we focus on the identification of cell type specific DEGs between sets of samples from different experimental conditions or genotypes, each measured at the single cell level, which has been identified as one of the grand challenges for single cell data analysis [17].

Expression quantitative trait locus (eQTL) [18-21] analysis is a special case of differential gene expression analysis where gene expression is combined with genetic information. A genetic variant associated with the transcription of a gene is called eQTL and allows for gaining insights into the molecular underpinnings of trait associated genetic variants. Using scRNA-seq, it is now possible to identify eQTLs in a cell type specific manner [22-25] and large scale efforts are currently underway [26]. In contrast to differential expression methods, linear regression models are typically used for the detection of eQTLs even in RNA-seq data sets [20,27], after transforming the count data to a normal distribution.

For the experimental design of transcriptome studies, researchers typically need to decide on sample sizes and technical parameters given certain constraints on resources. Constraints typically arise from the limited availability of samples or from the costs of the experiment. Power analysis should be used to make informed decisions about these parameters based on the statistical power to detect DEGs and eQTLs given certain assumptions about the expected effect sizes. The experimenter can determine either the power to detect DEGs and eQTLs, the required minimal sample size or the minimal effect size by keeping the other two parameters fixed based on prior knowledge or explicit assumptions. Power analysis is always tightly linked with the statistical testing procedure. Several methods have been established based on the theory of linear regression models [28] and the control of the false discovery rate [29-32] for microarray studies. For RNA-seq studies, power analysis methods based on the theory of negative binomial count regression [33,34], other parametric models [35-37], or simulations [38,39] have been proposed and benchmarked [40]. These power analysis methods can be used in combination with differential expression tools based on negative binomial count regression such as DESeq2 [5] or edgeR [41], which also perform well on scRNA-seq data [16,42-44]. Therefore, in principle methods for RNA-seq power analysis could also be applied to compute power or minimally required sample sizes for given effect sizes for single cell experiments.

To unlock RNA-seq power analysis methods for scRNA-seq data, the following aspects have to be addressed: 1) in interindividual comparisons, the variables of interest (e.g. genotype or phenotype) are measured at the individual level, whereas the expression levels are measured in many cells for each individual, and 2) scRNA-seq data is sparse, so only the most highly expressed genes are detectable. Here we address these aspects by formulating the identification of cell type specific DEGs and eQTLs as a (negative binomial) regression on ‘pseudo-bulk’ counts. This is allowing for the application of established power analysis methods. These are combined with a model for the probability of detecting cell type specific gene expression as a function of the number of cells of a certain type and the number of reads sequenced per cell. This allows for answering additional experimental design questions specific to single cell assays: 1) how many cells per individual are required? And 2) how deep should each cell be sequenced?

Previously, several individual aspects of single cell experimental design have been addressed. The comparison of sensitivity and accuracy of different technology platforms [45-47] has led to recommendations of sequencing depth. The minimal number of cells sequenced to observe a rare cell type with a given probability can be modelled with a negative binomial distribution [48,49] or multinomial distribution [50]. PowsimR [44] is a simulation based tool allowing for power analysis for the detection of DEGs between different cell types. First recommendations for the experimental design of interindividual comparisons with single cell resolution are currently being developed [51] in the context of eQTL analyses. However, the general question of interindividual comparisons between samples has not been addressed by these approaches. Moreover, handling more complex designs is not readily accessible for simulation based methods, but can be achieved with analytical power analysis methods.

Here we provide a unified resource for experimental design considerations of interindividual comparisons including the power to detect rare cell types as well as the power to detect DEGs and eQTLs. We derive data driven priors on expression distributions from single cell atlases of three different tissues, two from published studies [52,53] and a newly generated data set. We combine these with cell type specific priors for effect sizes based on DEGs and eQTL genes from bulk RNA-seq experiments on cells sorted by fluorescence activated cell sorting (FACS). The four DE studies [54-57] and one eQTL study [58] cover different biological applications to diseases such as asthma and cancer and to ageing. Together, this will enable researchers to design experiments across tissues with realistic effect size estimates. Our model provides the basis for rationally designing well powered experiments, increasing the number of true biological findings and reducing the number of false negatives. We provide our model and parameters as an open source R package *scPower* on github https://github.com/heiniglab/scPower. This also comprises a shiny app with a user-friendly graphical user interface, which is additionally available as a web server at http://scpower.helmholtz-muenchen.de/. All code to reproduce the figures of the paper is provided in the package vignette.

## Results

### Power analysis framework for scRNA-seq experimental design

For the power analyses we assume a gene x cell count matrix, based on UMI counts. Cells are annotated to an individual and a cell type or state. For the sake of simplicity, we will only consider discrete cell types / states. These can be derived by clustering and analysis of marker genes, potentially considering multiple levels of resolution [13] or using the metacell approach [59]. Individuals are annotated with different experimental factors. For the discussion we will consider a simple two group comparison, but more complicated experimental designs, which can be analysed with generalized linear models, can be treated analogously. To determine cell type specific differential expression between samples, gene expression estimates for each of N individuals (samples) and each cell type are required. We consider a ‘pseudo-bulk’ approach here, where cell type gene expression levels for each individual are approximated as the sum of UMI counts over all cells of the cell type. Three parameters determine the power and also the cost of a scRNA-seq experiment: 1) the number of samples *n*_*s*_, 2) the number of cells per sample *n*_*Ci*_ and 3) the number of reads sequenced per cell r. In order to determine the power of the experiment, we either need to make explicit assumptions or use prior knowledge about unknown experimental parameters. Figure 1 shows the dependency between different modifiable experimental parameters, unknown quantities and the expected outcomes. In our framework these dependencies are modelled as follows.

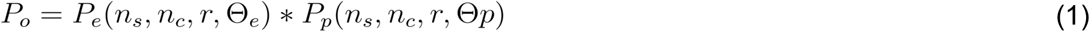

**Figure 1:**
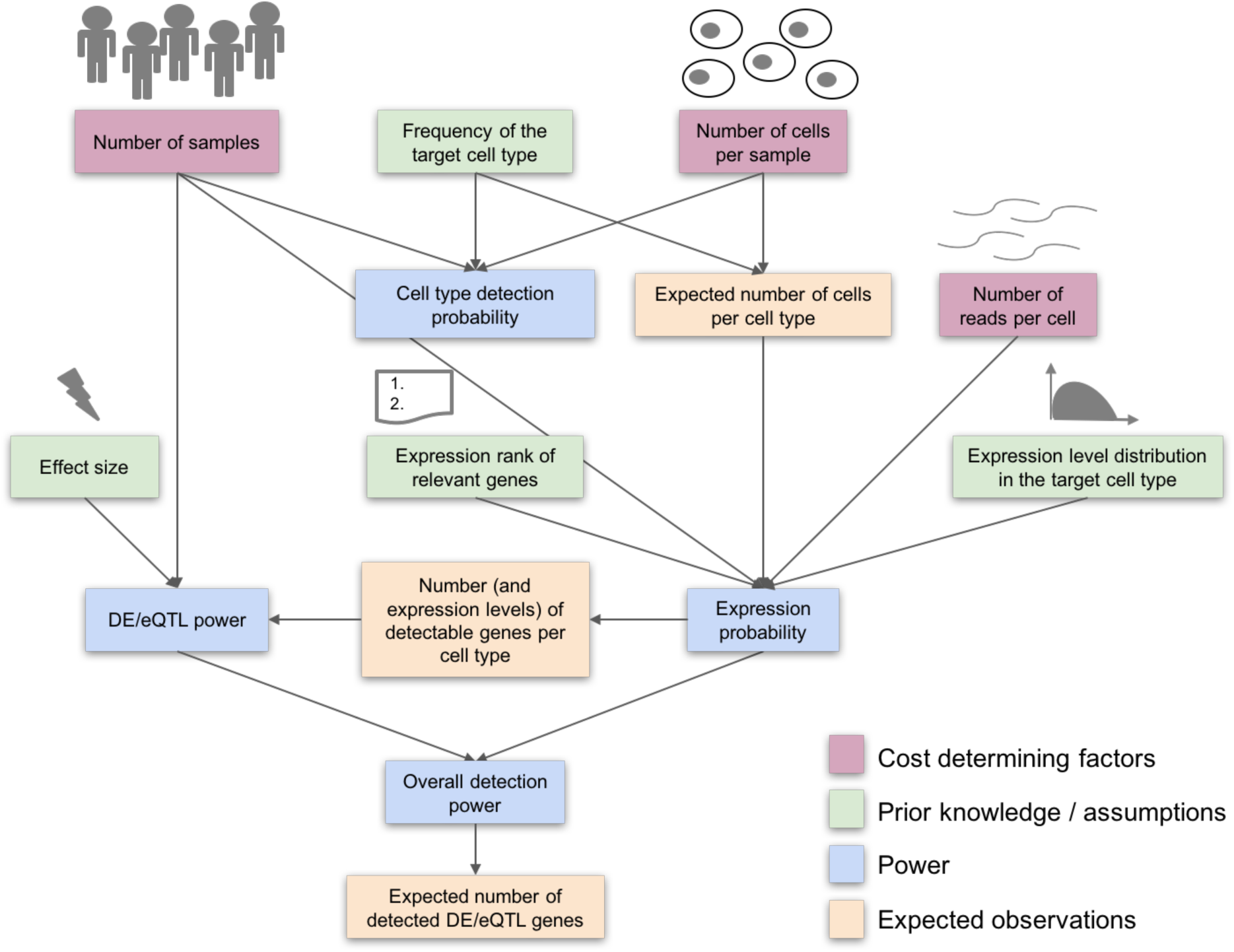
Dependence of experimental design parameters. The figure illustrates how the cost determining factors (purple: number of samples, number of cells per sample and number of reads per cell) are related to detection power (blue) and expected number of observations (orange). In addition, power and expected observations also depend on prior knowledge or assumptions (green).

#### Overall detection power P_o_

The expected number of detected DEGs depends on one hand on the probability to measure each of the relevant genes, based on their assumed expression levels. On the other hand it depends on the statistical power to detect DEGs of assumed effect sizes (see Methods, section <u>Overall detection power)</u>.

#### Expression probability P_e_

In scRNA-seq experiments individual cells are typically not sequenced to saturation, leading to sparse count matrices, where only highly expressed genes are detected with counts greater than zero. The overall number of transcripts as well as the number of transcripts of individual genes can be highly cell type specific [60]. We model the cell type specific gene detection probabilities depending on the number of reads sequenced per cell and on the number of cells of the cell type (see Methods, section <u>Expression probability model</u>).

#### DE / eQTL power P_p_

We use analytical power calculations based on linear or negative binomial regression models. The power of the statistical test for differential expression depends primarily on the number of samples and on the assumed effect sizes. In addition, the power of negative binomial regression models also depends on the expression level, with more power for highly expressed genes and less power for genes with high dispersion. The power also indirectly depends on the number of detectable genes, as this influences the multiple testing adjusted significance levels required in the power analysis. If expression quantitative trait loci are analysed instead of differential expression, the statistical power to detect eQTL genes is based on a linear model of transformed count data. Thus, in this model, the power depends only on the effect sizes and not on the individual gene expression levels (see Methods, section <u>Power analysis for differential expression</u> and <u>Power analysis for expression quantitative trait loci</u>).

#### Cell type detection probability

In exploratory analyses the goal is to observe as many cell types as possible. The power to observe rare cells depends on the (possibly unknown) frequency of this cell type and primarily on the number of cells sequenced per sample. Additionally, in the comparison between samples, it also depends on the total number of samples, as the rare cells need to be observed in all of the samples with certain minimal probability. We use negative binomial models to determine cell type detection probabilities (see Methods, section <u>Frequency of the rarest cell type</u>).

### scPower accurately models the number of detectable genes per cell type

In scRNA-seq experiments typically only highly expressed genes are detected with counts greater than zero [45-47]. The number of detectable genes per cell type depends on the number of reads sequenced per cell [46]. Here we consider the sum of all UMI counts per gene per cell type per individual as the gene expression measurement. Therefore, the number of detectable genes per cell type also depends on the number of cells of the cell type per individual. In the following sections, we specify a model for the number of detectable genes parameterized by the number of cells per cell type and individual and by the number of reads sequenced per cell. We fit the model using a scRNA-seq data set of PBMCs from 14 healthy individuals measured with 10X Genomics (Additional file 1: Figure S1, Table S1).

In our approach, we model UMI counts per gene in a particular cell type and individual as independent and identically distributed according to a negative binomial distribution. The mean of the distributions across all genes is modeled as a mixture distribution with a zero component and two left censored gamma distributions to cover highly expressed genes (see methods and Additional file 1: Figure S2). Subsampling the read depth of our data shows that the parameters of the mixture distribution are linearly dependent on the average count of unique molecular identifiers (UMI) (Additional file 1: Figure S3). Average UMI counts are related to the average number of reads mapped confidently to the transcriptome, which are in turn related to the number of reads sequenced per cell (Additional file 1: Figure S4). Taken together, we now have a model of per cell read counts across all genes parameterized by the number of reads sequenced.

Next, let us consider the count distributions of a particular gene in a particular cell type and individual. Based on the negative binomial distribution we can relate the distribution of UMI counts per cell to the distribution of the sum of UMI counts. We determine the expression probability for a gene using its expression rank to first compute the mean of the distribution of the sum of UMI counts based on the quantiles of our gamma mixture model and then use this distribution to compute the probability that the observed count is greater than a user defined minimal count threshold in at least a given number of individuals. Summing up these gene expression probabilities allows for modelling expected number of expressed genes. Figure 2 shows the number of expressed genes across cell types dependent on the number of cells of the cell type for varying read depth based on subsampling of our data. The observed numbers (solid lines) are closely matched by the expectation under our model (dashed lines) for genes with counts greater ten (Figure 2A) and with counts greater than zero (Figure 2B). While Figure 2A-B shows the results only for Run 5 of the PBMC data set, the fits of all runs can be found in Additional file 1: Figure S5,S6.

**Figure 2:**
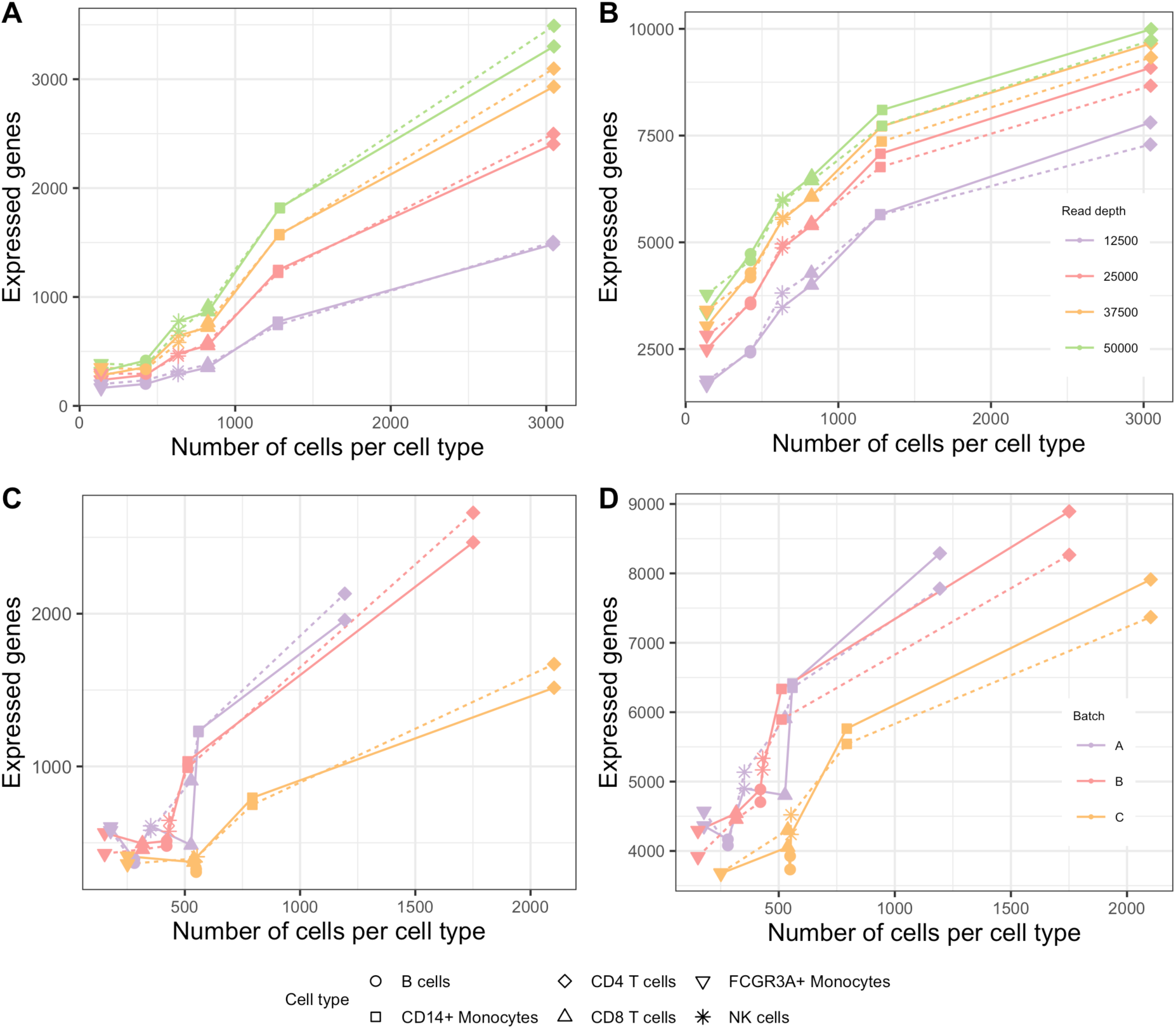
Expression probability model parameterized by UMI counts per cell. Plot A and B show the observed (solid) number of expressed genes and the number of expressed genes expected under our model (dashed) on the y-axis and the number of cells per cell type (cell type indicated by the point symbol) on the x-axis for Run 5 of the PBMC data set (Additional file 1: Table S1). The data is subsampled to different read depths (indicated by the color). Similarly, plot C and D show expressed genes for the three batches of the Ye data set. The definition for an expressed gene is based on a flexible user defined cutoff. Here it was parameterized: in Figure A and C, a gene is called expressed with count > 10 in more than 50% of the individuals, in Figure B and D, count > 0 in more than 50% of the individuals.

To validate our model, we applied it on a second PBMC data set [22] that was not used during parameter estimation (Figure 2C-D). The validation data set was measured at smaller read depth of 25,000 reads per cell and for a different sample size (batch A and B with 4 individuals and batch C with 8 individuals). The observed numbers are closely matched by the expectation under our model, which demonstrates that it can generalize well between data sets and different experimental parameters. Taken together, we have now a general model for the expected number of expressed genes, which is parameterized by the number of cells per cell type, the number of reads per cell. Of note, gene expression distributions are cell type specific and the model parameters have to be fitted from suitable pilot experiments, for instance from the human cell atlas project [61].

### scPower models the power to detect differentially expressed genes and expression quantitative trait genes

Negative binomial regression is a powerful approach for DEG analysis of both RNA-seq and scRNA-seq [16,42,43,62] and well tested tools such as DESeq [5,63] or EdgeR [41] are available. Here we use analytical methods for the power analysis of negative binomial regression models [64]. These power calculations are exact when analysing the data with models based on negative binomial regression, but DEG analyses with other tools might lead to different results. Power to detect an effect of a given effect size (log fold change) depends on the sample size, on the mean expression level and on the significance threshold alpha. The large number of parallel tests performed in a DEG analysis requires an adjustment of the significance level to avoid large numbers of false positive results. This can be achieved by controlling the family-wise error rate (FWER: probability of at least one false positive) using the Bonferroni method [65]: *a =* 0.05 / number of genes to guarantee FWER < 0.05. To obtain a range of typical effect sizes and mean expression distributions in specific immune cell types, we analysed several DEG studies based on FACS sorted bulk RNA-seq [54,55]. Combining our model for gene expression in scRNA-seq experiments with this power analysis of DE genes, we can calculate the overall detection power of DE genes in a scRNA-seq experiment as the product of the expression probability of the gene and the DE power for the gene (see Formula 1). To determine the expression probability for a gene, we use its expression rank from prior experiments to first compute the probability that the gene has more than a minimal number of counts in a specific cell type in at least a given number of individuals, as described above. Gene specific overall power is then derived based on the gene specific expression probability and the power to detect the gene as a DE gene based on fold changes from prior DEG studies.

Figure 3A shows that the overall detection power reaches up to 74% for fold changes from a study comparing CLL subtypes iCLL vs mCLL [54], when using 3,000 measured cells per cell type and individual and a total balanced sample size of 20, i.e. 10 individuals per group. The original study had a sample size of 6 individuals and detected 84 DEGs with median absolute log fold change of 2.8. For this parameter combination and prior, the DE power would reach even 96% for all DE genes of the study, however, only 74% are likely to be expressed. Overall, the DE power increases with higher number of measured cells and higher sample sizes, while the expression probability is mainly influenced by the number of measured cells. The influence of the sample size is not so pronounced in this example due to the small sample size of the reference study, for other reference studies it is more visible (Additional file 1: Figure S10). Notably, increasing the number of measured cells per individual and increasing the sample size both result in higher costs. In the next section, we show how a restricted budget affects the decision on the best parameter combinations to maximize the detection power. Especially, an increase in the sample size can generate high additional costs.

**Figure 3:**
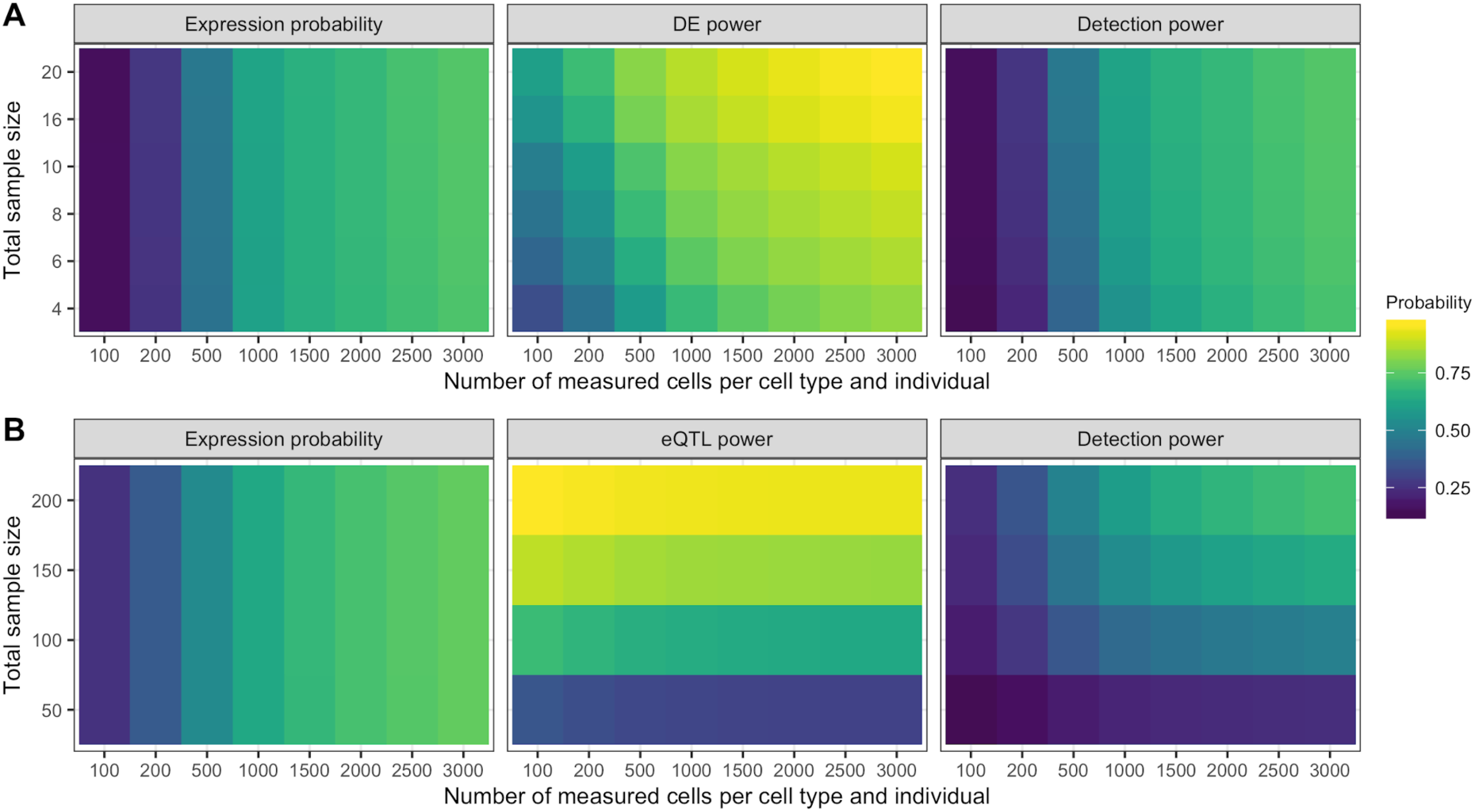
Expression probability, DE/eQTL power and overall detection power. Power estimation using data driven priors for A. DE genes and B. eQTL genes dependent on the total sample size and the number of measured cells per cell type. The detection power is the product of the expression probability and the power to detect the genes as DE or eQTL genes, respectively. The fold change for DEGs and the R^2^ for eQTL genes were taken from published studies, together with the expression rank of the genes. For A, the Blueprint CLL study with comparison iCLL vs mCLL was used, for B, the Blueprint T cell study. The expression profile and expression probabilities in a single cell experiment with a specific number of samples and measured cells was estimated using our gamma mixed models, setting the definition for expressed to > 10 counts in more than 50% of the individuals.

Similar detection ranges are found for the comparison of other CLL subtypes in the same study, while the detection power in a study of systemic sclerosis vs control were much lower with values up to 26% (Additional file 1: Figure S10). Smaller absolute fold changes in this study decrease the DE power and therefore also the overall detection power.

As a second application scenario, we analysed expression quantitative trait locus (eQTL) studies of specific immune cells [58]. The analysis of T cells in the study had a sample size of 192 and identified 5132 eQTL genes with a median absolute beta value for the strongest associated SNP of 0.89. Due to the very large number of statistical tests (~millions), simple linear models are usually applied to transformed read count data, as they can be computed very efficiently. Therefore, power calculations here are based on linear models [28] and the power is independent of the mean expression level.

Overall detection power for eQTL genes (Figure 3B) is more restricted by the expression probability of the genes than the eQTL power. The eQTL power increases with larger sample sizes, but decreases slightly with larger numbers of measured cells per individual and cell type. This is due to increased expression probability, which leads to an increased overall detection power, but also to a lower significance threshold due to the FWER correction (dividing by the number of expressed genes). This slightly reduces the eQTL power (Figure 3B). The same effect occurs for the DE power, but a higher number of measured cells generate higher mean expression values in the pseudo bulk data, which increase the power and counteract the negative effect of a more stringent significance threshold. In contrast, the eQTL power is modelled independently of the mean expression, so a higher number of genes has a slightly negative effect on eQTL power, but not on overall detection power. A maximal detection power of 68% and 69% was found for a sample size of 200 individuals and 3,000 measured cells per cell type and individual in the Blueprint eQTL data sets (Additional file 1: Figure S10).

### scPower optimizes the experimental parameters to maximize the detection power for a given budget

With this model for power estimation in DE and eQTL single cell studies, we are now able to optimize the experimental design for a fixed budget. The overall cost function for a 10X Genomics experiment is the sum of the library preparation cost and the sequencing cost (see Methods). The library preparation cost is defined by the number of measured samples and the number of measured cells per sample, while the sequencing cost is defined by the number of sequenced reads, which depends also on the target read depth per cell. Additional file 1: Figures S11, S12 show the three parameters maximizing detection power, given a fixed total budget. Figure 4 shows the optimization with expression priors from our PBMC data set, measured with 10X Genomics.

**Figure 4:**
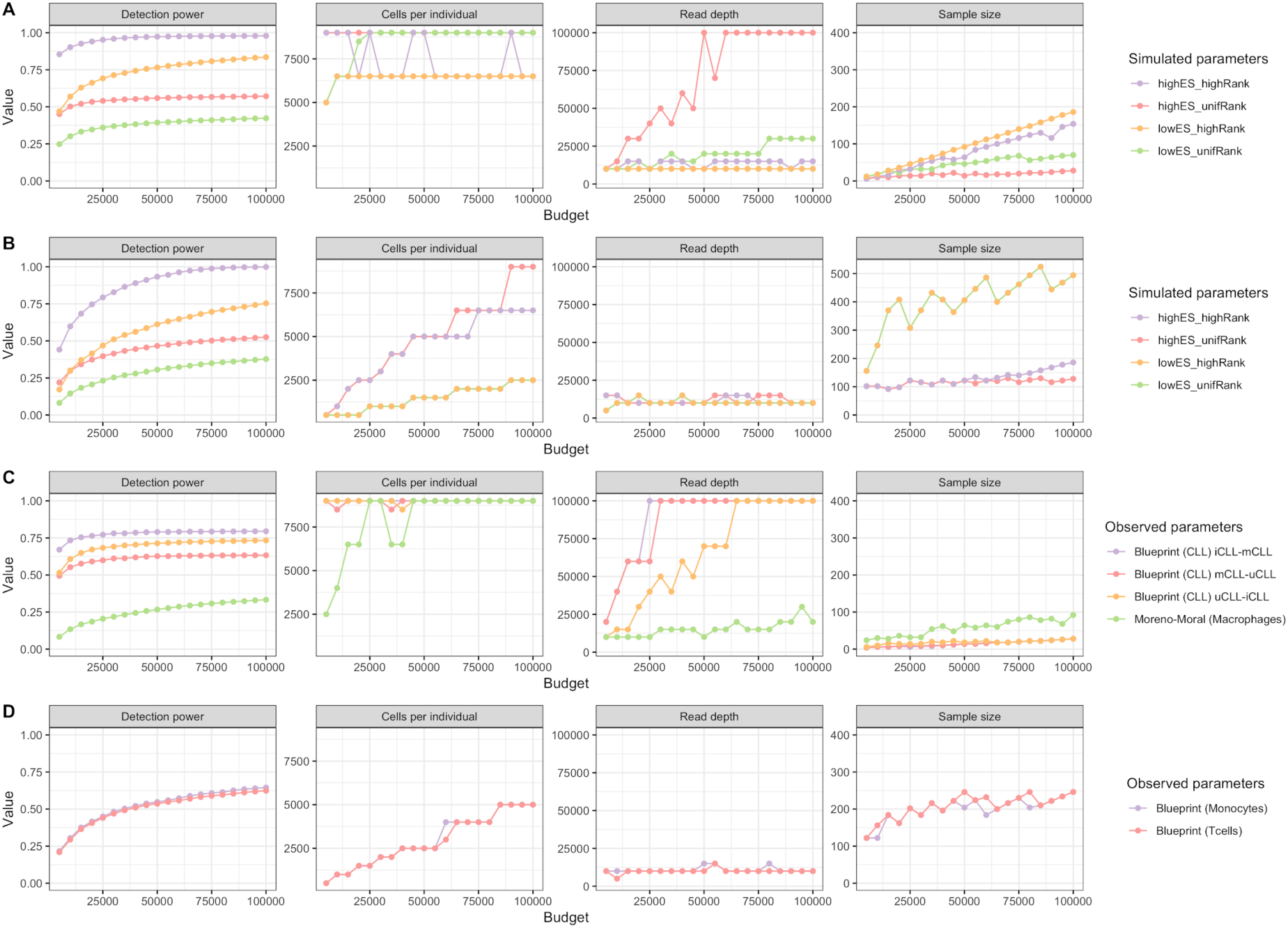
Optimal parameters for varying budgets and 10X Genomics data. The figure shows the maximal reachable detection power (y-axis, first column) for a given experimental budget (x-axis) and the corresponding optimal parameter combinations for that budget (y-axis, second till fourth column). The colored lines indicate different effect sizes and gene expression rank distributions. Subplots A-B visualize different simulated effect sizes and rank distributions (simulation names see text) for DEG studies (A) and eQTL studies (B) with models fitted on 10X PBMC data. Subplots C-D visualize effect sizes and rank distributions observed in cell type sorted bulk RNA-seq DEG studies (C) and eQTL studies (D) with model fits analogously to A-B.

We systematically investigated the evolution of optimal parameters for increasing budgets in four artificial scenarios for DEG (Figure 4A) and eQTL analysis (Figure 4B), four scenarios based on prior DEG (Figure 4C) and two scenarios on prior eQTL (Figure 4D) experiments on FACS sorted cells. The artificial scenarios reflect combinations of effect sizes (high, low) and expression levels (high, low) of DEGs and eQTL genes. We observed that the number of cells per individual is the major determinant of power, as this is the variable that is either directly set to maximum values or increased first in the optimization (Figure 4). This effect is least pronounced in the artificial eQTL scenario (Figure 4B), where small effect sizes require large sample sizes. For most DEG scenarios, the number of reads per cell is increased before increasing the sample size (Figure 4A,C), indicating that strong effects can be detected with relatively few samples, while the detection of expression required deeper sequencing. For eQTL scenarios first increasing the sample size is more beneficial than increasing the read depth (Figure 4B,D), which remains relatively low (10,000 reads per cell).

In the cost optimization, we also took into account that increasing the number of cells per lane leads to higher numbers of doublets, droplets with two instead of one cell, which need to be excluded from the analysis [22]. However, doublet detection methods such as Demuxlet [22] and Scrublet [66] enable faithful detection of those. We validated the doublet detection and donor identification of Demuxlet using our PBMC data set by comparing the expression of sex specific genes with the sex of the assigned donor (Additional file 1: Figure S1B) and found high concordance after doublet removal, also for run 5, which was overloaded with 25000 cells. The increase of the doublet rate through overloading was modeled using experimental data [67]. However, we observe in our own data set as well as in published studies [22,23] slightly higher doublet rates. Therefore, we consider the modeled doublet rate as a lower bound estimation. With a high detection rate of doublets, overloading of lanes is highly beneficial, since larger numbers of cells per individual lead to an increase in detection power, while not causing additional library preparation costs. Although, overloading leads to a decreasing number of usable cells and a decreasing read depth of the singlets, as doublets contain more reads, the overall detection power still rises strongly for both DE and eQTL studies.

### scPower generalizes across tissues and scRNAseq technologies

Our power analysis framework is applicable on data sets for other tissues besides PBMCS and for other single cell technologies besides 10X Genomics. We demonstrate this with a lung cell data set measured by Drop-seq [53] and a pancreas data set measured by Smart-seq2 [52]. The model of the expression probability needs to be adapted slightly for other technologies, while the DE/eQTL power calculation remains the same as for 10X.

Smart-seq2 is a plate-based technology, generating read counts from full-length transcripts. Therefore, we express the count threshold for an expressed gene relative to one kilobase of the transcript. We fitted the expression model including the transcript length in the size normalization factor of the count model. Additionally, we modelled the doublet rate as a constant factor. In contrast, Drop-seq is a droplet-based technology similar to 10X Genomics and exactly the same model can be used. However, as we are lacking the experimental data to fit an appropriate model for overloading, we set the doublet rate again constant. With these adaptations our expression probability model works well for other tissues and technologies (Additional file 1: Figure S13).

Analogously to Figure 4, the evolution of parameters for simulated priors (Figure 5A,B) and observed priors (Figure 5C,D) was evaluated across the other technologies. Similar trends are observed for the Drop-seq lung data as for the 10X PBMC data set in the artificial scenarios (Figure 5A) as well as for observed priors from cell type sorted bulk studies (Figure 5C). In both cases, the number of cells per individual is the major determinant of power. Overall, lower power is observed for the Smart-seq2 pancreas study (Figure 5B,D). In contrast to 10X and Drop-seq, the optimal number of reads per cell is much higher and the number of cells per individual and sample size is increased only at higher budgets for both the artificial and data driven priors.

**Figure 5:**
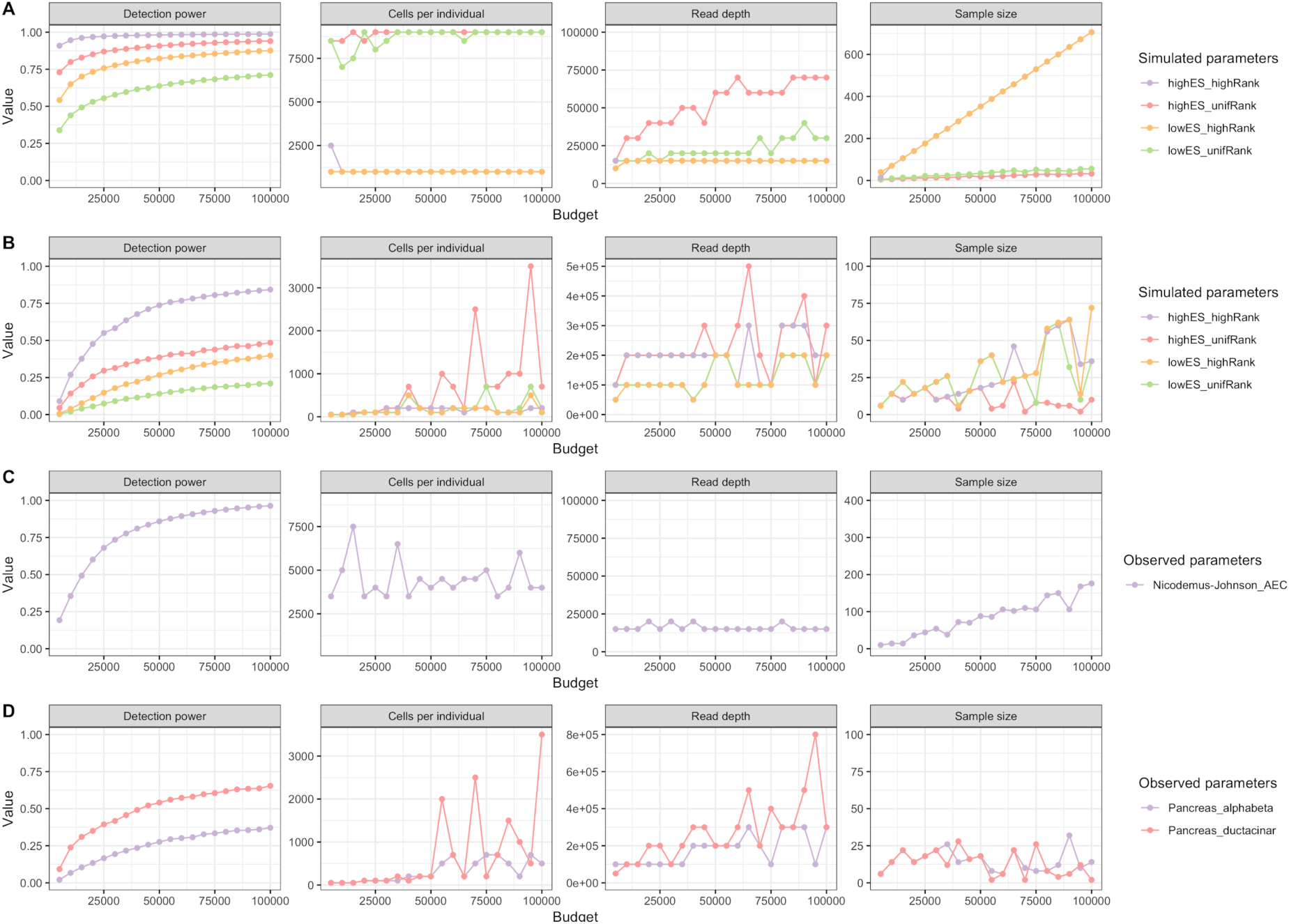
Optimal parameters for varying budgets and Drop-seq and Smart-seq2 data. The figure shows the maximal reachable detection power (y-axis, first column) for a given experimental budget (x-axis) and the corresponding optimal parameter combinations for that budget (y-axis, second till fourth column). The colored lines indicate different effect sizes and gene expression rank distributions. Subplots A-B visualize different simulated effect sizes and rank distributions (simulation names see text) for DE studies with models fitted on Drop-seq lung data (A) and Smart-seq2 pancreas data (B). Subplots C-D visualize effect sizes and rank distributions observed in cell type sorted bulk RNA-seq DE studies with model fits analogously to A-B.

### Power to detect rare cell types

In exploratory analyses, the goal is to observe as many cell types as possible. The power to observe rare cells depends on the frequency of this cell type, the number of cells sequenced per individual and the total number of individuals. Following [48], we model this problem using the negative binomial distribution (see methods). Here we demonstrate the approach using prior knowledge of cell proportions in PBMCs from the literature to determine the number of cells required for each individual to detect a minimal number of cells of a specific type. The rarest immune cell type we considered are dendritic cells, which occur in PBMCs with a frequency of 1.5%. Consequently, more than 1000 cells per individual are required to observe at least ten dendritic cells in all individuals with probability greater than 95%, while only about 300 cells are required for NK cells, which have a frequency of 7% in PBMCs (Additional file 1: Figure S14). The comparison for varying numbers of individuals shows that the number of cells required for each individual is most strongly affected by the frequency of the cell type and only to a smaller degree by the number of individuals.

## Discussion

We have introduced *scPower*, a method for experimental design and power analysis for interindividual differential gene expression and eQTL analysis with single cell resolution. Based on realistic data driven from multiple tissues and artificial priors, we have observed that the number of cells is a major determinant of power in droplet based assays, followed by sample size and read depth.

The number of cells drives power by increasing the sensitivity of gene expression detection. Previous analyses such as [46] have recommended 1 Mio reads per cell when comprehensive gene expression detection is desired. Our analyses suggest that aggregating shallowly sequenced transcriptomes of a large number of cells of the same cell type is a more cost efficient way than increasing read depth to increase the sensitivity for individual level gene expression analysis. Most likely, multiple independent library preparations in individual cells lead to an improved sampling of the transcriptome as compared to fewer independent libraries sequenced more deeply, an effect that has previously been analysed in the context of variant detection [68]. The number of cells to be sequenced has previously been considered with respect to the power of detecting rare cell types [48,49], however, its effect on gene expression sensitivity is equally important.

Optimal read depths (∼10000) are relatively low compared to previous recommendations [45,47,69,70]. In a systematic analysis of ERCC spike in expression it has been shown that the accuracy of the measurements is not strongly dependent on the sequencing depth and consistently high (∼0.9 Pearson correlation for 10X and Drop-seq) for a read depth of 10,000 reads per cell [46]. Hence, we expect accurate individual level gene expression quantification with the optimized experimental design.

The number of cells and sequencing depth also determine the accuracy of the extraction of gene expression programs, which are critical for the annotation of cell types [71]. Shallow sequencing of higher number of cells has achieved equal accuracy as deeper sequencing of fewer cells [71]. In line with our findings, it has thus been recommended to shallowly sequence more cells [71].

As expected, the sample size is mostly dependent on the effect size, with low effect sizes requiring large sample sizes and consequently optimal setting with high sample size typically lead to low sequencing depth and relatively low number of cells.

Previous experimental design methods for scRNA-seq like powsimR [44,47] allow for the simulation of scRNA-seq read counts on the single cell level and model the fold changes between groups of cells. Here we consider the fold changes between groups of individuals. To enable the usage of the simulation model of powsimR for the comparison of individuals, a model relating the effect sizes observed on the individual level to single cells would be required. This is particularly challenging especially for continuous individual level covariates. The pseudo-bulk approach presented here allows for leveraging well established power analysis methods based on (generalized) linear models. While it represents a baseline method for power analysis, however, it has a number of limitations. First, the negative binomial regression model for pseudo-bulk inspired by DESeq [5,63] might not be the most powerful method for assessing individual level differential expression. Our current framework is tightly linked to this approach and cannot easily be extended to arbitrary analysis methods, this is however the case for all analytical power analysis methods. Second, it requires a discrete cell type definition. Therefore, continuous cell annotations such as pseudo time would need to be discretized before the power analysis. Third, our method works on raw data and is not able to process imputed gene expression matrices. Imputation is a popular approach to maximize the number of quantifiable genes by borrowing information of gene expression profiles of cells within the same cell type or even between cell types [72-75]. Imputation within the same cell type will not assign expression values greater than zero to genes that were zero in all cells of the cell type. Thus the number of expressed genes (count > 0) in the pseudo bulk approach should be very similar to within cell type imputation. Some imputation methods such as [73,74] do model the negative binomial means per gene, which we use in our model, so these results could in principle be integrated in our model. Fourth, we did not address the power for the detection of variance QTLs from scRNAseq data [24] due to the lack of data driven priors for the effect sizes.

Several practical considerations should be addressed when using our approach. First, our data driven priors only allow for reliably assessing the overall power in sample sizes that are smaller or roughly equal to the sample size of the pilot data sets from which the effect sizes were estimated. Consequently, a larger sample size will identify new significant DEGs with lower effect sizes, which were not identified in the smaller pilot study and thus not included in the computation of the overall detection power. Second, our current modeling of the doublet rate using reference values from 10X Genomics is a lower bound compared to the doublet rates we estimate for our own data and to rates reported by other studies [22,23]. Thus, actual experiments might result in higher doublet rates and lower number of usable cells. Last, the choice of a threshold on the number of reads required for a gene to be called expressed influences also choice of optimal parameters. Here we used a threshold of >10 reads, however, some eQTL analyses of bulk RNAseq data advocate using >0 reads [27], whereas DESeq2 automatically chooses the threshold that optimizes the number of discoveries at a given FDR by applying the independent filtering strategy [5,76]. Best practice guidelines for differential gene expression with RNA-seq recommend cutoffs that remove between 19%-33% of lowly expressed genes, depending on the analysis pipeline [77]. These percentages correspond to 1-10 reads per million sequenced, which translates to 1-5 UMI counts for a median of around 5000 UMI counts per cell in our data set. Our gene expression probability model is cell type specific and has to be fitted based on realistic pilot data. We have shown that our model can be applied to 10X Genomics, Drop-seq and Smart-seq2 and we would expect that it is applicable also to other technology platforms.

Importantly, experimental design recommendations here are optimized for differential expression between individuals. Other applications might result in very different optimal experimental designs. For instance, co-expression analysis requires a high number of quantified genes per cell, especially when one is interested in cell type specific co-expression and comparison of such co-expression relations between individuals. Furthermore, the power to identify new rare cell types by clustering analysis of scRNA-seq data might have different optimal parameters [49].

The human cell atlas project has outlined a ‘skydive’ strategy of iteratively sampling human cells with increasing resolution to build a reference map of healthy human cells [61,78]. In combination with the human cell atlas reference transcriptomes and cell type annotations *scPower* will provide the foundation for building a comprehensive resource for the experimental design of interindividual gene expression comparisons with cell type resolution across all organs systems covered in the human cell atlas.

## Conclusions

*scPower* is a unified resource for experimental design considerations of interindividual comparisons with cell type resolution. It models the power to detect rare cell types as well as the power to detect DEGs and eQTLs. Based on data driven priors on expression distributions from single cell atlases of three different tissues and cell type specific priors for effect sizes based on DEGs and eQTLs from bulk RNA-seq experiments, we show that shallow sequencing of high numbers of cells per individual lead to higher overall power than deep sequencing of fewer cells. Our model generalizes across different tissues and scRNAseq technologies. The method is implemented in an R package with a user friendly graphical user interface and is freely available on github. Our model will provide the basis for rationally designing well powered experiments, increasing the number of true biological findings and reducing the number of false negatives.

## Methods

### Collection of PBMCs

Blood was collected from psychiatric control individuals according to the clinical trial protocol of the Biological Classification of Mental Disorders study (BeCOME; ClinicalTrials.gov TRN: NCT03984084) at the Max Planck Institute of Psychiatry. All individuals gave informed consent. Perinuclear blood cells (PBMCs) were isolated and cryopreserved in RPMI 1640 medium (Sigma-Aldrich) supplemented with 10% Dimethyl Sulfoxide at a concentration of roughly 1M cells per ml.

### Single cell RNA-sequencing

For single-cell experiments, 14 cell vials from different individuals (7 male and 7 female) were snap-thawed in a 37°C water bath and serially diluted in RPMI 1640 medium (Sigma-Aldrich) supplemented with 10% Fetal Bovine Serum (Sigma-Aldrich) medium. Cells were counted and equal cell numbers per individual were pooled in two pools of 7 individuals each. Cell pools were concentrated and resuspended in PBS supplemented with 0.04 % bovine serum albumin, and loaded separately or as a combined pool with cells of all 14 individuals on the Chromium microfluidic system (10X Genomics) aiming for 8,000 or 25,000 cells per run. Single cell libraries were generated using the Chromium Single Cell 3’library and gel bead kit v2 (PN #120237) from 10X Genomics. The cells were sequenced with a targeted depth of approximately 50,000 reads per cell on the HiSeq4000 (Illumina) with 150 bp paired-end sequencing of read2 (exact numbers for each run in Additional file 1: Table S1).

### Preprocessing of the single cell RNA-seq data

We mapped the single cell RNA-seq reads using CellRanger version 2.0.0 and 2.1.1 [79]. Demuxlet was used to identify doublets and to assign cells to the correct donors [22]. Additionally, Scrublet version 0.1 was run with a doublet threshold of 0.28 to identify also doublets from cells which originate from the same donor [80]. Afterwards, the derived gene count matrices from CellRanger were loaded into Scanpy version 1.4 [81]. Cells identified as doublets or ambivalent by Demuxlet and Scrublet were removed, as well as cells with less than 200 genes or more than 2,500 genes and with more than 10% counts from mitochondrial genes.

### Verification of Demuxlet assignment using sex errors

We validated the donor assignment and doublet detection of Demuxlet by testing if assigned cells express sex specific genes correctly. Xist expression was taken as evidence for a female cell, expression of genes on the Y chromosome as evidence for a male cell.

The male sex error shows the fraction of cells assigned to a male donor among all cells expressing Xist (count > 0). The threshold for the female error was set less strictly, as mismapping of a few reads to the chromosome Y occurs also in female cells. Instead, the female sex error indicates which fraction of cells is assigned to a female donor among all cells having more reads mapped to chromosome Y than the *frac*_*female*_ quantile of all cells, with *frac*_*female*_ being the overall fraction of cells assigned to a female donor among all cells. TPM mapped to chromosome Y is calculated by counting all reads mapped to chromosome Y, excluding reads mapped to the pseudoautosomal regions, times 10^6^ divided by the total number of read counts per cell.

Both error rates are calculated twice, once with all cells and once without doublets from Demuxlet and Scrublet.

### Cell type identification

We performed the cell type identification according to the Scanpy PBMC tutorial [82]. Genes which occurred in less than 3 cells were removed. Counts were normalized per cell and logarithmized. Afterwards the highly variable genes were identified, the effect of counts and mitochondrial percentage regressed out. We calculated a nearest neighbour graph between the cells, taking the first 40 PCs, and then clustered the cells with a Louvain clustering [83]. Cell types were assigned to the clusters using marker genes (Additional file 1: Table S2).

### Influence of read depths

We used subsampling to estimate the dependence of gene expression probabilities on read depths. The fastq files of all 6 runs were subsampled using fastq-sample from fastq-tools version 0.8 [84]. The number of reads was downsampled to 75%, 50% and 25% of the original number of reads. CellRanger was used to generate count matrices from the subsampled reads. Donor, doublet and cell type annotation were always taken from the full runs with all reads.

### Expression probability model

The gene expression distribution of each cell type was modeled separately because there are deviations in RNA content between different cell types [60]. The UMI counts *x* per gene across the cells of a cell type are modeled by a negative binomial distribution. We used DESeq [63] to perform the library size normalization as well as the estimation of the negative binomial parameters. The standard library size normalization of DEseq and the variant “poscounts” of DESeq2 [5] were both used, depending on the quality of the fit for the specific data set. For the PBMC 10X data set (Additional file: Table 1), the standard normalization was taken, for the Drop-seq lung and the Smart-seq2 pancreas datasets the poscount normalization, which is more suitable for sparse data. Only cell types with at least 50 cells were analysed to get a robust estimation of the parameters.

The negative binomial distribution is defined by the probability of success *p* and the number of successes *r*:

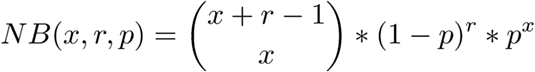

DESeq uses a parametrization based on mean *μ* = *P r/*(1 ^—^ *P)* and dispersion parameter *disp* = l/r.

We formulated the definition of an expressed gene in a flexible way so that users can adapt the thresholds. In general, a gene is called *expressed* in a cell type if the sum of counts *V* over all cells of the cell type within an individual is greater than *n* in more than *k* percent of the individuals. We assume a negative binomial distribution for the counts *x* of each gene in each cell type c with *μc* and *disp*_*C*,_ omitting indices of gene and individual for clarity. The sum of the gene counts *y* over all cells *n*.*cellsCtIndiv* of an individual in the cell type follow a negative binomial distribution with *μ*_*sum*_ *= n*_*cellsCtIndiv*_ * *μe* and *disp*_*surn*_ *= disp*_*c*_*/n*_*cellsCtIndiv*._ The probability that the sum of counts in one individual is greater than *n* is:

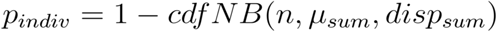

To define a gene as expressed, we require that it can be found in a certain fraction of more than *k* percent in all *n*_*s*_ individuals. The expression probability of a gene is obtained from a binomial distribution:

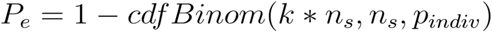

So in total, the expected value of the expected number of expressed genes can be defined as

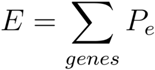

To generalize the expression probability model also for unseen data sets, the distribution of mean values Me over all genes in a cell type is modelled as a mixture distribution with three components, a zero component and two left-censored gamma distributions:

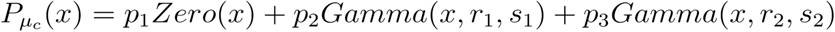

The model is an adaption of the distribution used in the single cell simulation tool Splatter [85]. The largest part of the mean values can be fitted with one gamma distribution, a small fraction with high expressed gene outlier with the second gamma distribution. The genes with zero mean values originate from two sources. Either, the gene is not expressed or the expression level is too low to be captured in the setting. The lower bound for the expression level at which both Gamma distributions are censored depends on the number of cells measured for this cell type *n*_*cellsCt*._ The smallest expression level to be captured is 1/*n* _*cellsCt*._ The density of the gamma distribution is parametrized by rate *r* and shape *s:*

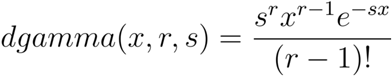

For modeling of the gamma parameters, also the parameterization by mean *μ* and standard deviation *sd* is used:

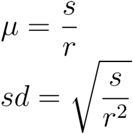

The relationship between the mean UMI counts per cell and the gamma parameters (mean and standard deviation of the two gamma distributions) is linear and *β* values are estimated by linear regression. The mixture proportion of the zero component *P*_*1*_ is linearly decreasing with the mean UMI counts, also estimated by linear regression. The lower bound of *P*_*1*_ is set to a small positive number: 0.01. In contrast, the mixture proportion of the second gamma component *P*_*3*_ is modelled as a constant, independent of the mean UMI counts. We set it to the median value of all fits per cell type. The mixture proportion of the first gamma component *P*^*2*^ is 1 ^—^ *P*_*1*_ ^—^ *P*_*3*_ and is linearly increasing with increasing mean UMI counts.

The number of transcriptome mapped reads is linearly related to the logarithm of the mean UMI counts per cell, with an increasing read depth leading to a saturation of UMIs. 10X Genomics describes this also with the sequence saturation parameter. The exact logarithmic saturation curve depends on multiple biological and technical factors, therefore, it needs to be fitted for each experiment individually. However, *scPower* provides example fits from the different scenarios observed in our analysis.

The dispersion parameter is estimated dependent on the mean value using the dispersion function fitted by DESeq. The parameters of the mean-dispersion curve showed no correlation with the mean UMI counts, therefore the mean of the parameters of the dispersion function across all runs and subsampled runs were taken, resulting in one mean-dispersion function per cell type.

### Power analysis for differential expression

The power to detect differential expression is calculated analytically for the negative binomial model [64]. An implementation of the method can be found in the R package MKmisc. Parameters are sample size, fold change, significance threshold, the mean of the control group and the dispersion parameter (assuming the same dispersion for both groups). Zhu et al. implemented three different methods to estimate the dispersion parameter, we chose method 3 for the power calculation, which was shown to be more accurate in simulation studies in the paper. More complex experimental designs can be addressed using the method of [86].

### Power analysis for expression quantitative trait loci

Additionally to the DE analyses, the use of scRNA-seq for the detection of expression quantitative trait loci (eQTLs) was evaluated. Here, the power to detect an eQTL is calculated using an F-test and depends on the sample size *n*_*s*_, the coefficient of determination *R*^*2*^ of the locus and the chosen significance threshold *a. R*^2^ is calculated for the pilot studies from the regression parameter *β*, its standard error *se* (*β*) and the sample size *N* of the pilot study:

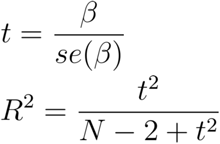

The implementation pwr.f2.test of the R package pwr is used for the F-test [28]. The degrees of freedom of the numerator are one and of the denominator are *n*_*s*_ − 2, the effect size is *f*2 = *R*^*2*^*/(* 1 - *R*^*2*^)

### Overall detection power

Assuming independence between the expression probability and power to detect DEGs / eQTLs, the overall detection power for DEGs is the product of the two probabilities. Expression probabilities were determined based on the gene expression rank in the observed (pilot) data. The number of considered genes *G* was set to 21,000, the number of genes used for fitting of the curves. Ranks i were transformed to the quantiles 1/*G* of the gamma mixture model parameterized by the mean UMI counts to obtain the mean *μ*_*c*_ of the negative binomial model, which is in turn used to compute the expression probability.

To quantify the overall power of an experimental setup, we compute the expected fraction of detected DEG / eQTL genes with prior expression levels and effect sizes derived from the pilot data. We obtain gene expression ranks of DEGs / eQTLs and their corresponding fold changes, to compute overall detection power for each gene. DE power is computed using a threshold *a* that is controlling the family-wise error rate (probability of at least one false positive among *E* expressed genes) as 0 05/*E* For eQTLs we were accounting for the number of genes that we test, assuming that there is a maximum of one cis-eQTL per gene. The overall power of the experimental setup is then the average detection power over all prior DEG/eQTL genes.

### Pilot data sets

Realistic DE and eQTL priors, i.e. effect sizes and expression ranks, were taken from sorted bulk RNA-seq studies of matching tissues (PBMCs, lung and pancreas). For all studies, the significance cut-off of the DE and eQTL genes was set to FDR < 0.05 and the expression levels of the genes were taken from FPKM normalized values. When published, we took directly the effect sizes, otherwise we recalculated the DE analysis with DEseq2.

#### Differential gene expression

To get realistic estimates for effect sizes (fold changes), data sets from FACS sorted bulk RNA-seq studies were taken [54,55]. The data sets were used to rank the expression level of the DEGs among all other genes using the FPKM values. The cell types used in the studies were matched to our annotated cell types in PBMCs for the expression profiles. The expression profile of CD14+ Monocytes was used for the study of Macrophages, the profile of CD4+ T cells for the CLL study.

Lung cell type specific priors were obtained from a DE study of freshly isolated airway epithelial cells of asthma patients and healthy controls [56]. As no effect sizes were reported, the analysis was redone with the given count matrix from GEO using DEseq2.

A DE study analyzing age-dependent gene regulation in human pancreas [57] was used to get pancreas cell type specific priors. We obtained expression ranks and gene length, which is needed for proper normalization of Smart-seq2 expression values.

#### eQTLs

We used eQTL effect and sample sizes from the Blueprint study on bulk RNA-seq of FACS sorted Monocytes and T cells [58]. Neutrophils were excluded as they are no PBMCs. We took the most significant eQTL for each gene, using a significance cutoff of 10^−6^. We compared the FPKM normalized expression levels of the eQTL genes among all other genes to get the expression rank for each eQTL gene. Effect sizes were derived from the slope parameters of the linear regression against genotype dosage, its standard error and the sample size of the study.

### Cost calculation and parameter optimization for a given budget

The overall experimental cost for a 10X Genomics experiment is the sum of the library preparation cost and the sequencing cost. It can be calculated dependent on the three cost determining parameters sample size ^*n*^*s*, number of cells per sample *n*_*c*_ and the read depth *r*. The library preparation cost is determined by the number of 10X kits, depending on how many samples are loaded per lane *n*_*sLane*_ and the cost of one kit *cost*_*kit*_ The cost of a flow cell *cost*_*fiowCell*_ and the number of reads per flow cell *r*_*fiowCell*_ determine the sequencing cost.

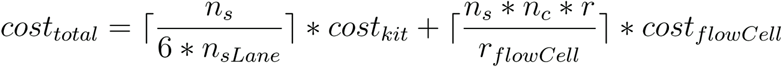

We optimized the three cost parameters for a fixed budget to maximize the detection power. A grid of values for number of cells per individual and for the read depth was tested, while the sample size is uniquely determined given the other two parameters and the fixed total costs. As an approximation of the sample size, the ceiling functions from the cost formula were removed.

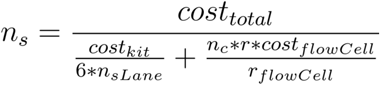

The same approach can also be used with a grid of sample size and cells per sample or read depth. In general, two parameters need to be chosen and the third parameter is uniquely determined given the other two and the fixed experimental cost.

Given the three cost parameters, the detection power for a specific cell type and a specific DE or eQTL study can be estimated. However, we also have to account for the appearance of doublets during the experiment. The fraction of doublets depends on the number of cells loaded on the lane. Following the approach of [48], we model the doublet rate *d* linear dependent on the number of recovered cells, using the values from the 10X User guide of V3 [67]. A factor of 7.67 * 10^−6^ was estimated, so that *d =* 7.67 * 10^−6^ * *n*_*c*_ * *n*_*sLane*_.

The number of usable cells per individual used for the calculation of detection power is then *n*_*usabieCells*_ *=* (1 — *d)* * *n*_*c*_ w_e_ assume that nearly all doublets are detectable using Demuxlet and Scrublet and that these cells will be discarded during the preprocessing of the data set. The expected number of cells for the target cell type with a frequency of *f* will be *f* * (1 - *d*) * *n*_*c*_

A second effect of doublets is that the read distribution is shifted, as doublets contain more reads than singlets. Again following the approach of [48], we assume that doublets contain 80% more reads than singlets. In the following, the ratio of reads in doublets compared to reads in singlets is called doublet factor *df*, a factor of *df =* 1.8 is assumed in the calculations in this manuscript. Therefore, depending on the number of doublets, the read depth of the singlets will be slightly lower than the target read depth.

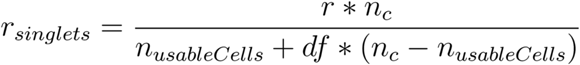

Additionally, the mapping efficiency is taken into account. Assuming a mapping efficiency of 80%, *r*_*mapped*_ *=* 0-8 * *r*_*singlets*_ mapped read depth remains. In the power calculation, the number of usable cells per cell type will be used instead of the number of cells and the mapped read depth instead of the target read depth.

Instead of defining the number of samples per lane directly, usually the number of cells loaded per lane *ncellsLane* is defined. So, the doublet rate per lane can be directly restricted. We use in our analyses *n*_*Cell*_*sLane* = 20,000, which leads to a doublet rate of at most 15.4%. The number of individuals per lane can be derived directly as *n*_*sLane*_ *= \n*_*cellsLane/n*c_].

### Simulation of effect sizes and gene rank distributions

Model priors, i.e. effect sizes and gene rank distributions, were derived from FACS sorted bulk RNA-seq to get realistic assumptions. Additionally, we simulated different extreme prior distributions to evaluate their influence on the optimal experimental parameters. The log fold changes for the DE studies were modeled as normally distributed. High effect size distributions were simulated with a mean of 2 and a standard deviation of 1, low effect sizes distributions with a mean of 0.5 and standard deviation of 1.

Effect sizes (*R*^2^ values) for the eQTL studies were obtained by sampling normally distributed Z scores and applying the inverse Fisher Z Transformation. Because very small values are not observed due to the significance threshold, the normal distribution is truncated to retain values above the mean. High effect sizes were simulated with a mean of 0.5 and standard deviation of 0.2, low effect sizes with a mean of 0.2 and a standard deviation of 0.2. A similar standard deviation was also observed in the pilot data.

250 DEGs were simulated and 2000 eQTL genes. The ranks were uniformly distributed, either over the first 10,000 genes or the first 20,000 genes. This leads to four simulation scenarios for each, high and low effect sizes (ES) and high or uniformly distributed expression ranks, called in the studies highES_highRank, lowES_highRank, highES_unifRank and lowES_unifRank.

### Evaluation of Drop-seq and Smart-seq2 data

We validated our expression probability model for other tissues and single cell RNA-seq technologies. Two data sets of the human cell atlas were used for that, a Drop-seq data set measured in lung tissue [53] and a Smart-seq2 data set measured in pancreas tissue [52].

The Drop-seq technology is also a droplet-based technique, similar to 10X Genomics. The same model can be used, only adapting the doublet and cost parameter. However, as there was no data available to model the linear increase of the doublet rate during overloading correctly, the doublet rate was modeled instead as a constant factor and the library preparation costs were estimated per cell. *scPower* provides models for both cases and with the necessary prior data, users can also model the overloading for Drop-seq.

Smart-seq2 is a plate-based technique, which produces full length transcripts and read counts instead of UMI counts. To compensate the gene length bias in the counts, the definition of an expressed gene was adapted to at least *n* counts per kilobase of transcript, resulting in a gene specific threshold of *n/geneLength* * 1000. The gamma mixed distribution of the mean gene expression levels is modelled using length normalized counts, but the gene length is required as a prior for the dispersion estimation and the power calculation, as DEseq uses counts, which are not normalized for gene length. These priors can be obtained together with the effect sizes and the expression ranks from the pilot bulk studies. In the simulation of non-DE genes, an average mean length of 5,000 bp is assumed. The linear relationship of the parameters of the mixture of gamma distributions is modeled directly based on the mean number of reads per cell. Doublets also appear in Smart-seq2, but as a constant factor, not increasing with a higher number of cells per individual. We observed for the parameter of the DEseq dispersion model a linear relationship with the read depth, which was not visible for Drop-seq and 10X Genomics. So, instead of taking the mean value per cell type, a linear fit is modeled for Smart-seq2.

For both data sets, the cell type frequencies varied greatly among individuals, therefore an estimation of expressed genes in a certain fraction of individuals could not be validated, as this requires similar cell type frequencies for each donor. Instead, the expressed genes were estimated to be above a certain count threshold in all cells of a cell type, independent of the individual.

Both data sets were subsampled to investigate the effect of the read depth. The Drop-seq reads are subsampled using fastq-tools version 0.8 [84] and the subsampled UMI count matrix was generated following the pipeline previously described in [87]. The Smart-seq2 read matrix was subsampled directly using the function *downsampleMatrix* of the package *DropletUtils* [88].

We compared the budget restricted power to our PBMC 10X Genomics results, using the same simulated effect sizes and distribution ranks as well as matched observed priors from FACS sorted bulk studies.

### Frequency of the rarest cell type

The probability to detect at least *n*_*cellsCtIndiv*_ cells of a specific cell type in each individual depends on the frequency of the cell type /, the number of cells per individual *n*_*c*_ and the number of individuals *n*_*s*_. For one individual, the minimal number of cells can be modeled using a cumulative negative binomial distribution [48]

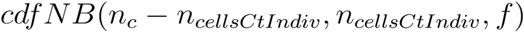

and for all individuals as

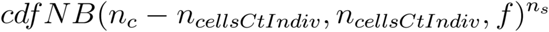

The cell type frequencies were obtained by literature research, the frequencies in PBMC are approximately twice as high as in whole blood [89]. All other parameters can be freely chosen (dependent on the expected study design).

## Declarations

### Ethics approval and consent to participate

All investigations have been carried out in accordance with the Declaration of Helsinki, including written informed consent of all participants. Study conduct complies with the recommendations by the ethics committee of the Bavarian Chamber of Physicians, Munich and the ethics committee of the Ludwig-Maximilian University, Munich. Applicable national and EU law, in particular the General Data Protection Regulation (GDPR) (Regulation (EU) 2016/679) has been followed.

Permission for using the data has been obtained from the Biobank of Max Planck Institute of Psychiatry. Consent for secondary use of the existing data has been obtained. In compliance with the consent for secondary use, the data generated in this project will be stored and can be used for future research. All data has been pseudonymized.

### Consent for publication

Written informed consent of all participants allows for publication of data in an access controlled repository.

### Availability of data and materials

The datasets generated and/or analysed during the current study are available in the European Genome Phenome Archive (EGA), accession number pending.

## Authors’ contributions

KTS and MH conceived the power analysis framework and analyzed the data. MH, FJT, EBB and HL designed the scRNA-seq experiment. EBB planned the BeCOME study and recruited the study participants. CC and AB generated scRNA-seq data in PBMCs. KTS and MH wrote the manuscript with input from all authors. All authors approved the final manuscript.

## Acknowledgements

We thank Thomas Walzthoeni for bioinformatics support provided at the Bioinformatics Core Facility, Institute of Computational Biology, Helmholtz Zentrum München. We thank Elisabeth Graf and Thomas Schwarzmayr for help in sequencing. We thank the BeCOME study team at the Max Planck Institute for Psychiatry, including the BioPrep core unit for their contribution to control individuals recruitment and characterizations, as well as collection of PBMCs. We thank Maren Büttner for insightful discussion and proofreading of the manuscript.

