## Supplementary Material for "Design and power analysis for multi-sample single cell genomics experiments"

#### Supplementary Figures

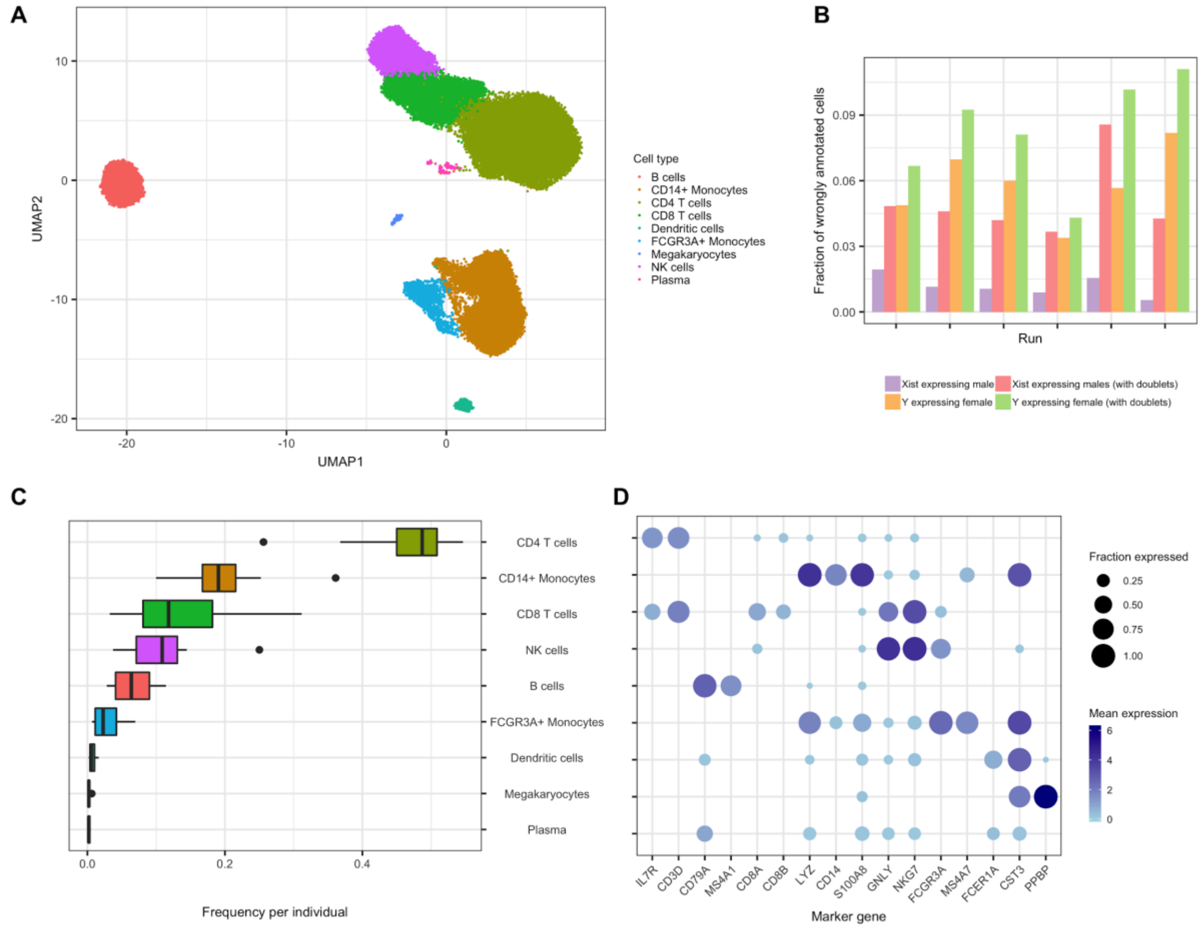


**Figure S1**: *PBMC data set*. A. UMAP visualization of all 6 runs, clustered using Louvain and annotated to cell types using marker genes. B. Evaluation of Demuxlet assignment to the different individuals by testing the expression of sex specific genes per run. The error rate "Xist expressing male" shows which fraction of cells is assigned to a male donor from all cells expressing Xist. The "Y expressing female" shows which fraction of cells is assigned to a female donor from all cells having more reads mapped to chromosome Y than the median value. Both error rates decrease when Demuxlet and Scrublet doublets are removed. C. Cell type frequencies for each individual. D. Marker gene distribution over the Louvain clusters. The color of the point visualizes its mean expression in the cluster, the size of the dot in how many cells of the cluster it is expressed (expression level larger than 0).

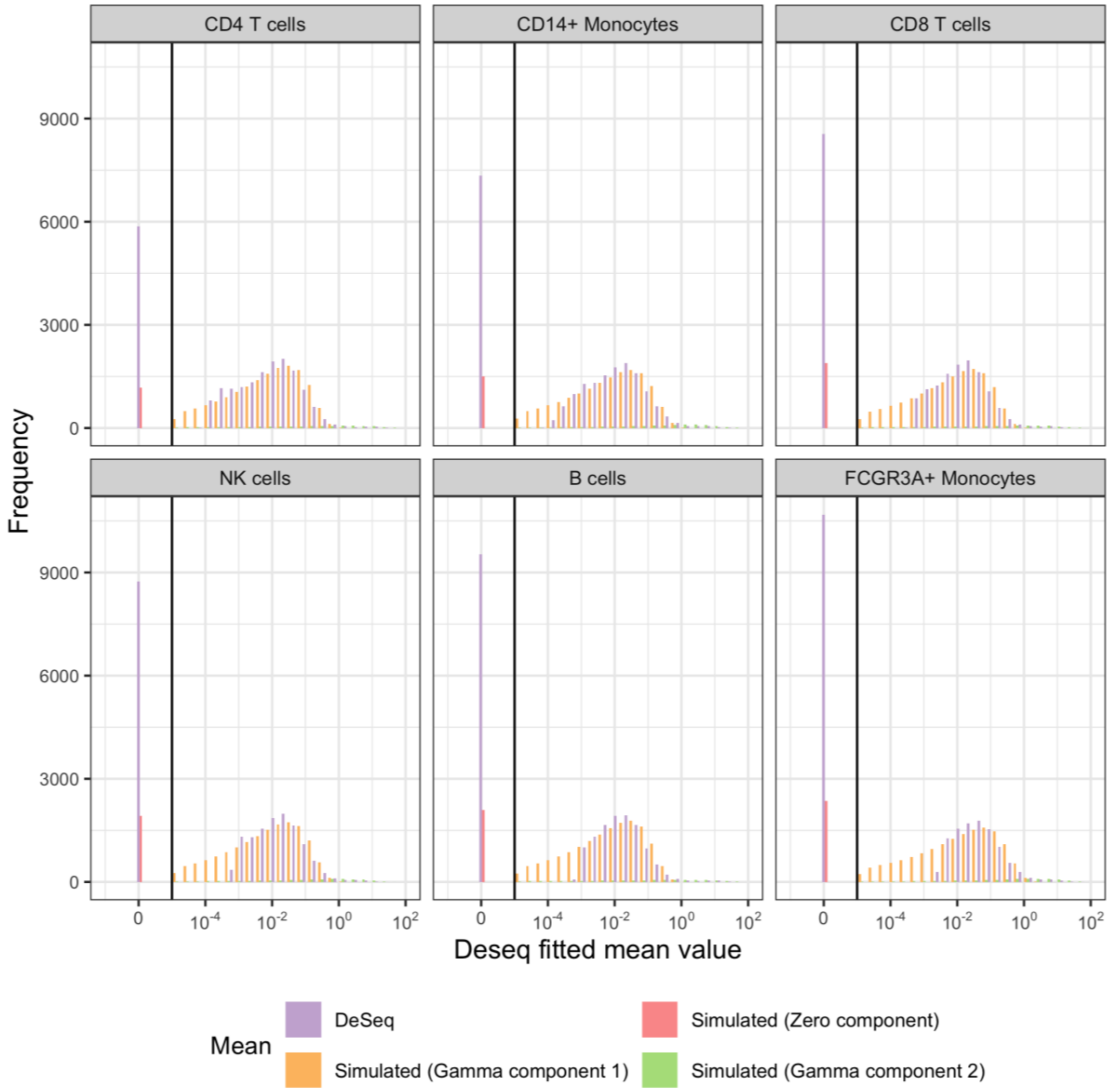


**Figure S2**: *Evaluation of gamma mixture fits for expression means*. Gamma mixed fit over all gene expression means for Run 5 of our PBMC data set per cell type. For each cell type, 21,000 genes are selected including all genes with counts > 0 in the cell type.

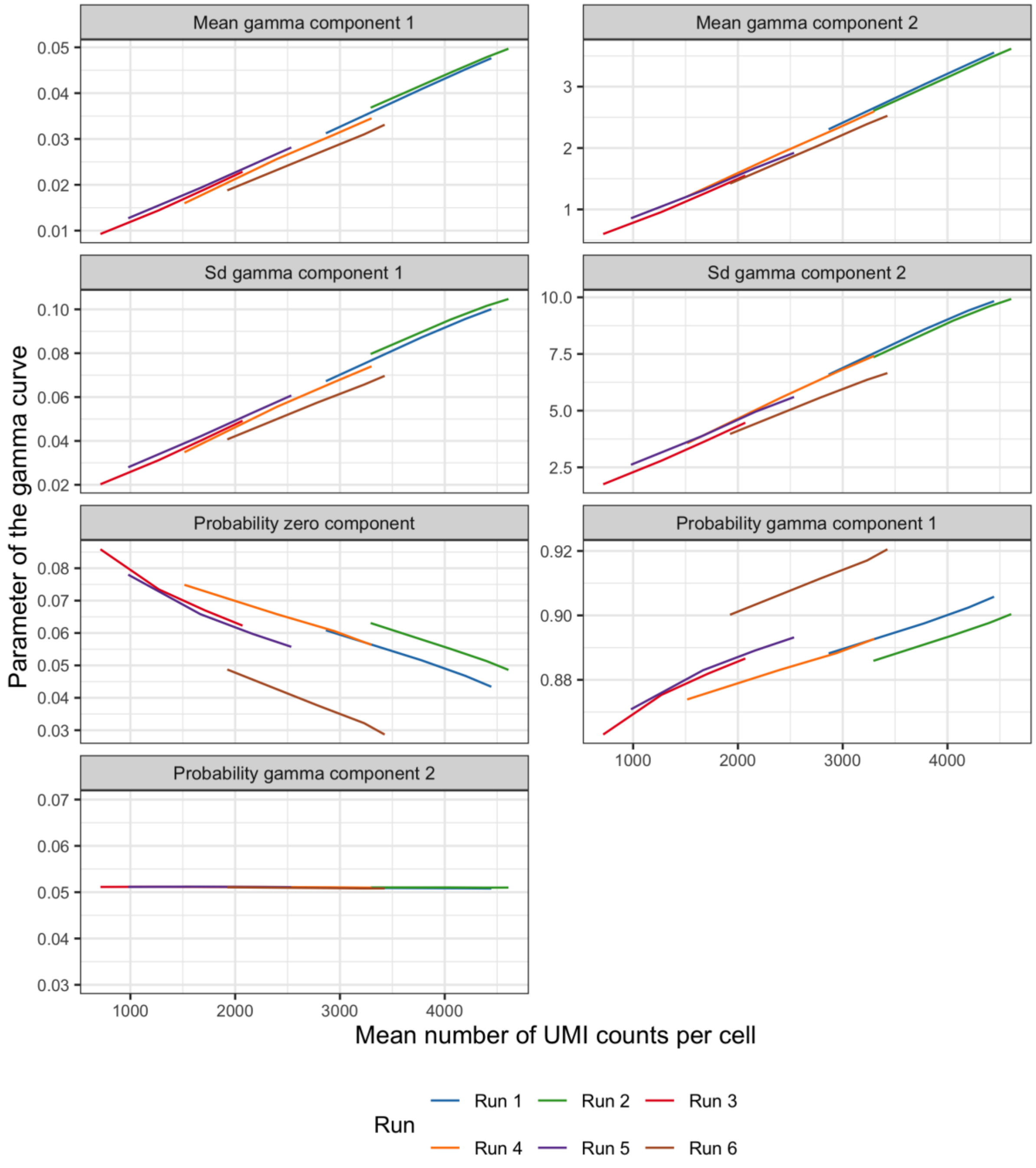


**Figure S3**: *Relationship between the parameters of the mixture fit and the mean number of UMI counts per cell.* The two left censored gamma distributions of the mixture are parametrized over their means and standard deviations, additionally there are three probability parameters showing the proportion of each of the three components, the zero component and the two gamma components. The fits were performed for each cell type separately, here shown for the CD4 T cells. There is a linear relationship between the mean and standard deviation parameters of the gamma components and the mean UMI counts. Also the probabilities of the zero component and the first gamma component show a linear relationship to the mean UMI counts. The probability parameter of the second gamma component stays constant. The other cell types show the same pattern.

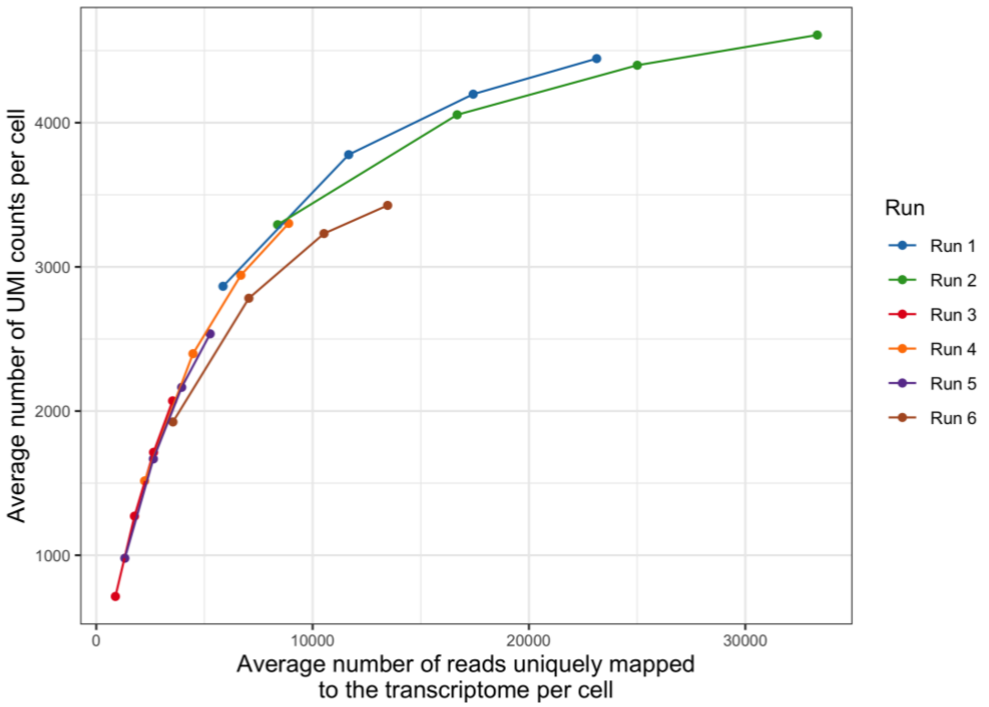


**Figure S4**: *Relationship between UMI counts per cell and average number of reads that were uniquely mapped to the transcriptome per cell.*

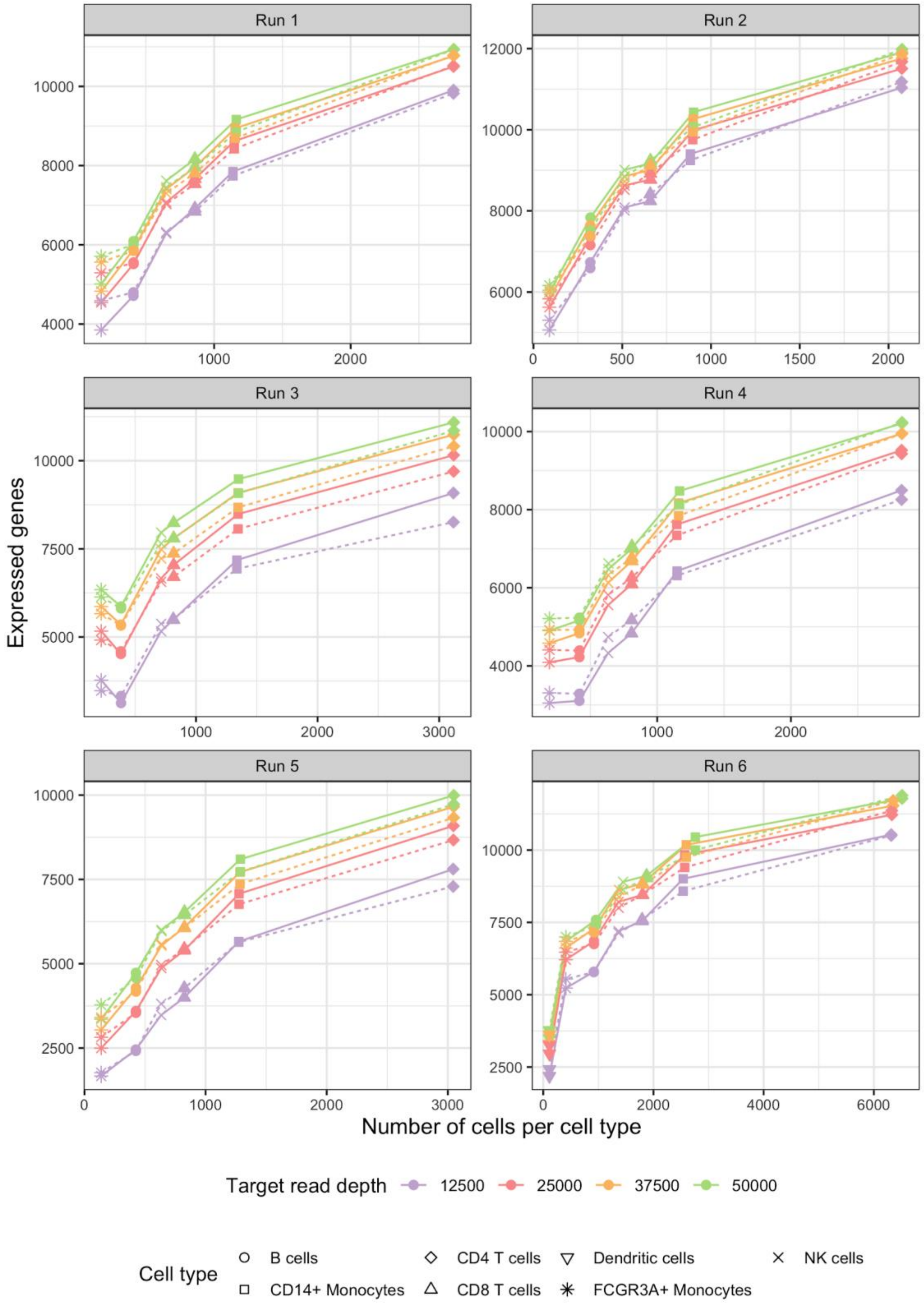


**Figure S5**: *Expression probability model for a count threshold larger than zero*. Gene curve fits using our gamma mixed models parameterized over UMI counts per cell for all our six runs. The definition for an expressed gene was parameterized in the way that a gene needs to have a count > 0 in more than 50% of the individuals.

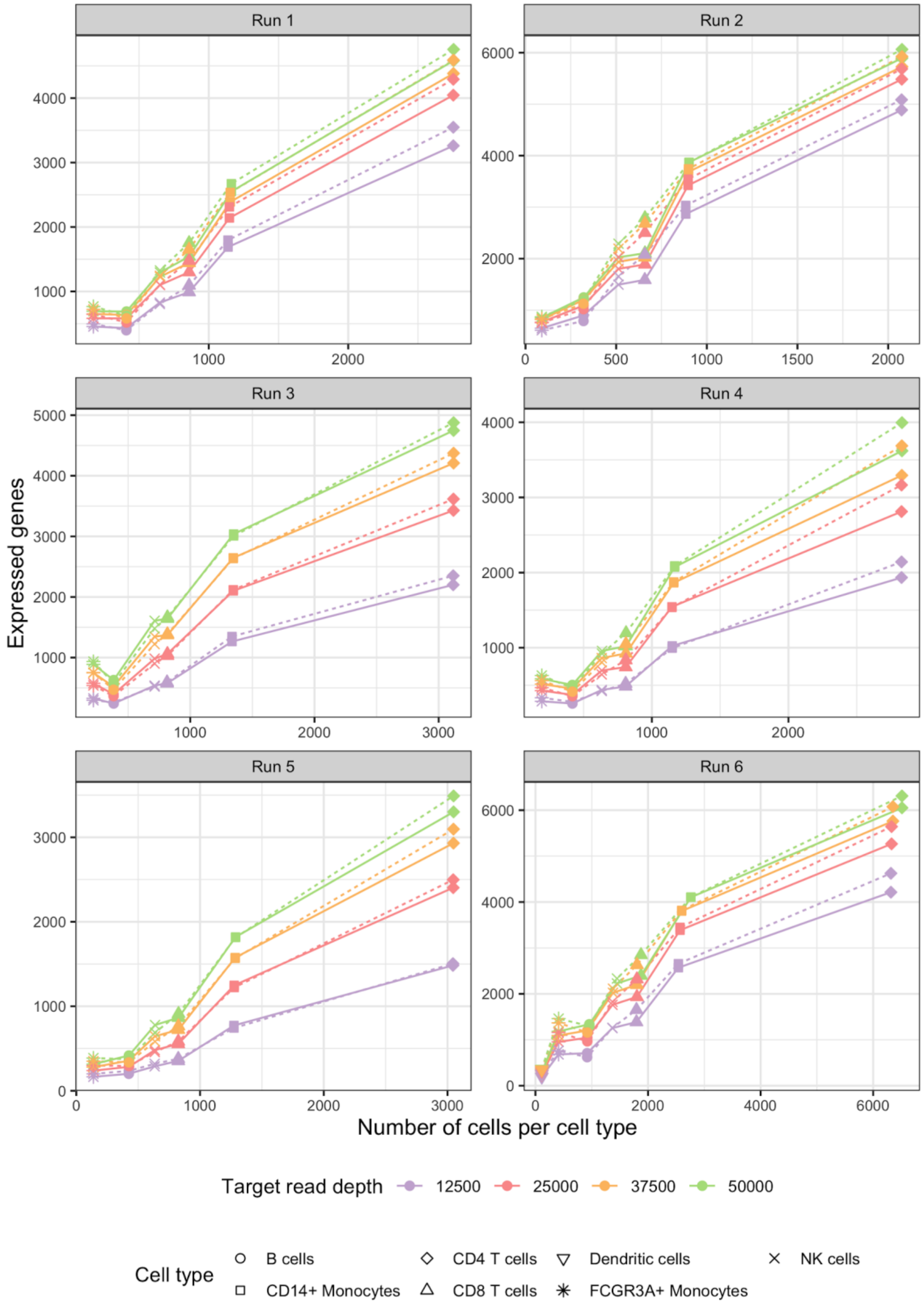


**Figure S6**: *Expression probability model for a count threshold larger than ten.* Gene curve fits using our gamma mixed models parameterized over UMI counts per cell for all our six runs. The definition for an expressed gene was parameterized in the way that a gene needs to have a count > 10 in more than 50% of the individuals.

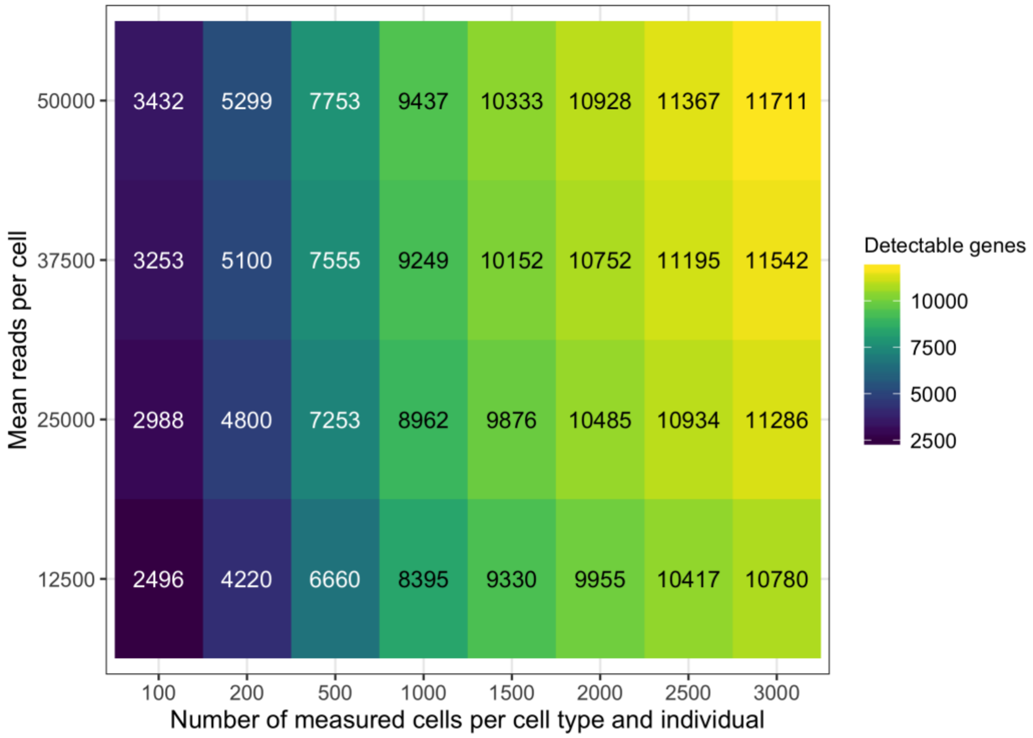


**Figure S7**: *Influence of the number of measured cells per cell type and individual and the mean reads per cell on the number of detectable genes*. Exemplarily, it is shown for the cell type CD4 T cells and a study size of 14 individuals. The definition for a detectable gene is set to at least 10 counts in more than 50% of all individuals.

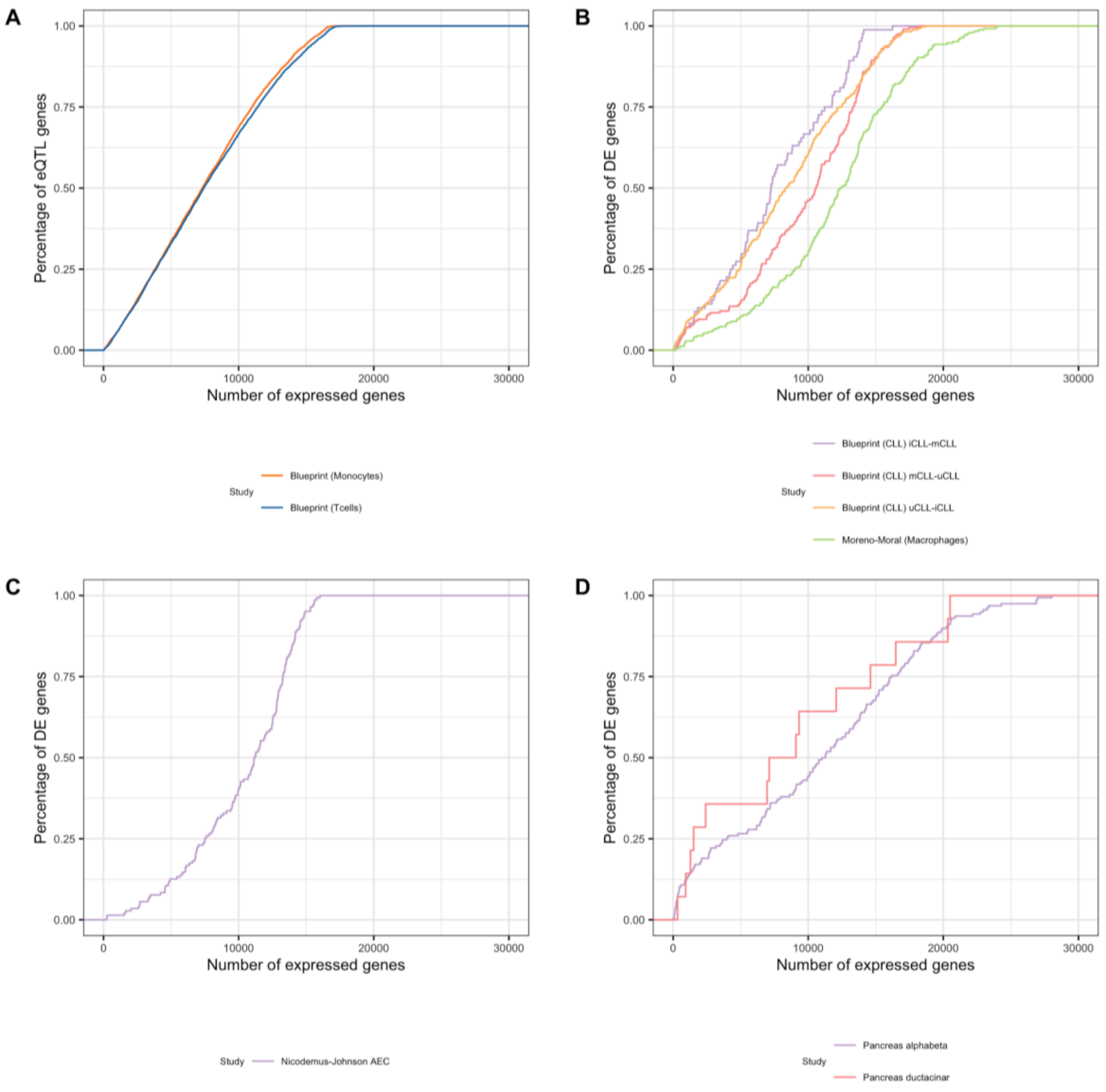


**Figure S8**: *Expression rank priors from cell type sorted bulk studies*. Gene expression distribution relative to all genes (i.e. gene expression ranks) gained from cell type sorted bulk studies for A. eQTL studies of PBMCs B. DE studies of PBMCs C. DE studies of lung tissue and D. DE studies of pancreas tissue. The figure shows which percentage of the DE/eQTL genes is expressed for a certain number of expressed genes.

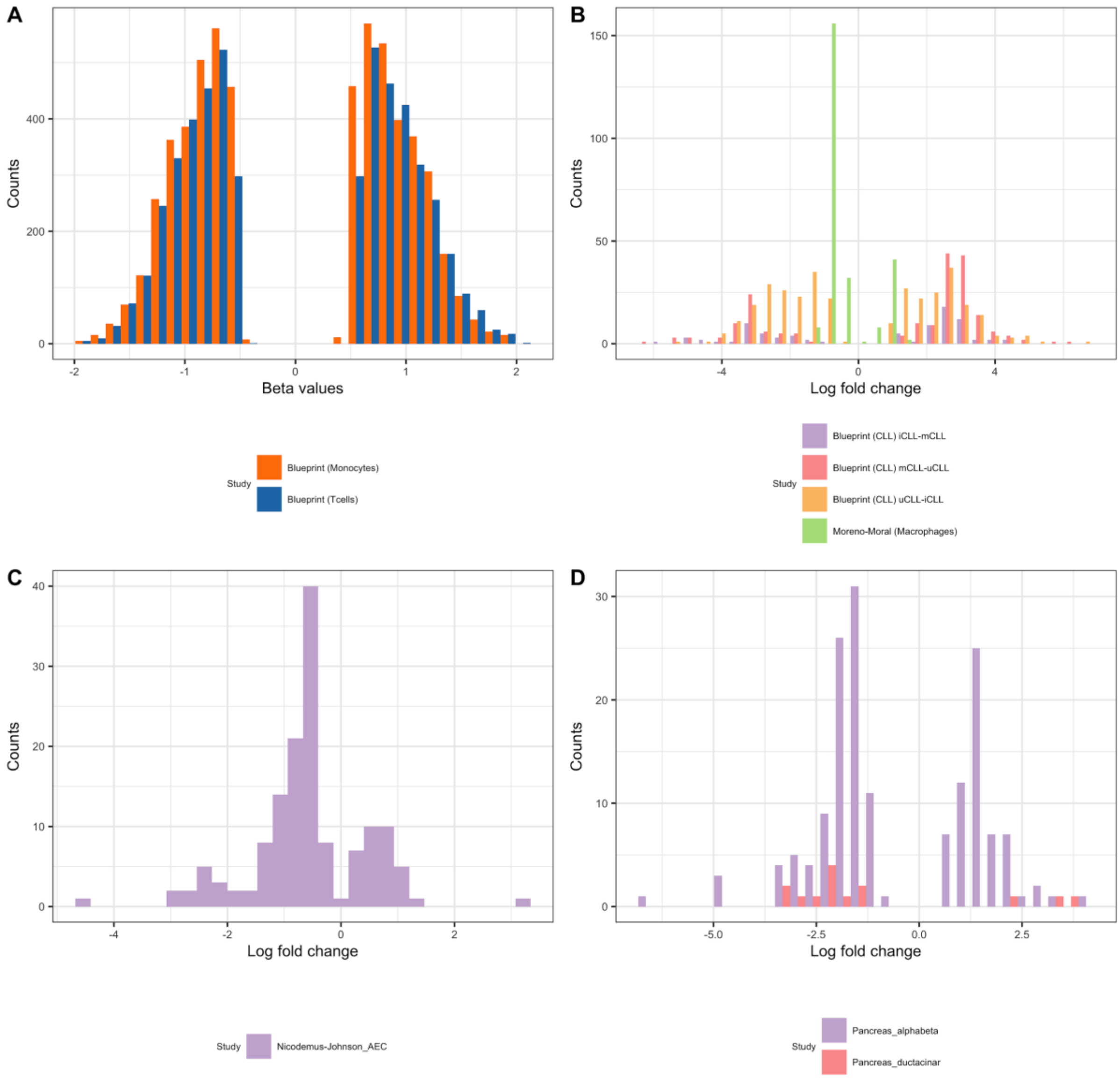


**Figure S9**: *Effect size priors from cell type sorted bulk studies*. Effect sizes gained from cell type sorted bulk studies for A. eQTL studies of PBMCs B. DE studies of PBMCs C. DE studies of lung tissue and D. DE studies of pancreas tissue. The effect size is quantified as beta values for eQTL studies and as log fold changes for DE studies.

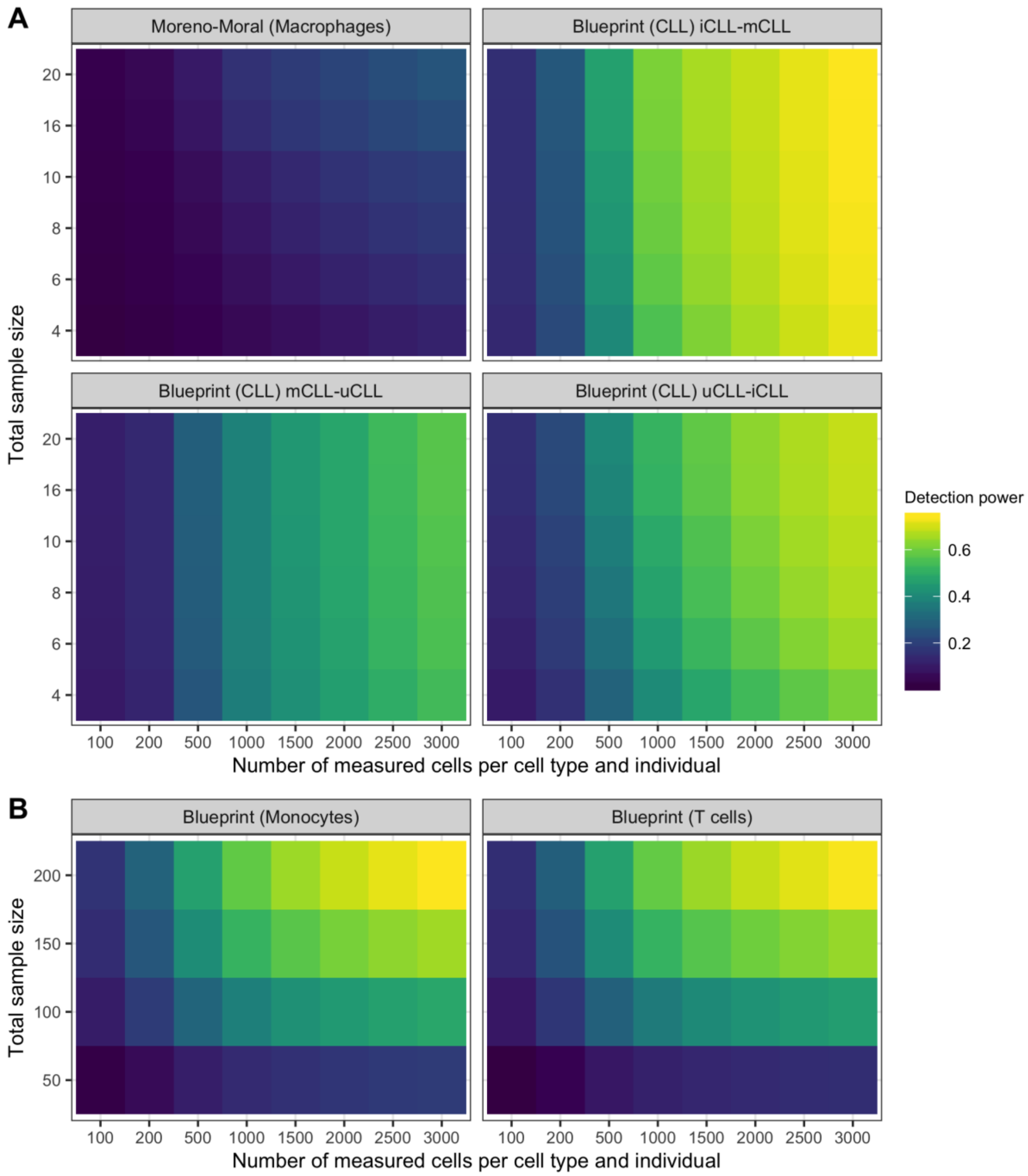


**Figure S10**: *Detection power using observed priors from reference studies*. Detection power for A. DE genes and B. eQTL genes dependent on the study, total sample size and the number of measured cells per cell type for a transcriptome mapped read depth per cell of 20,000. The detection power is the product of the probability that the gene is expressed and the power to detect it as a DE or eQTL gene, respectively, assuming that it is expressed. The fold change for DE genes and the R^2 for eQTL genes is taken from the published study, together with the expression rank of the genes. The expression profile in a single cell experiment with a specific number of samples and measured cells is estimated using our gamma mixed models.

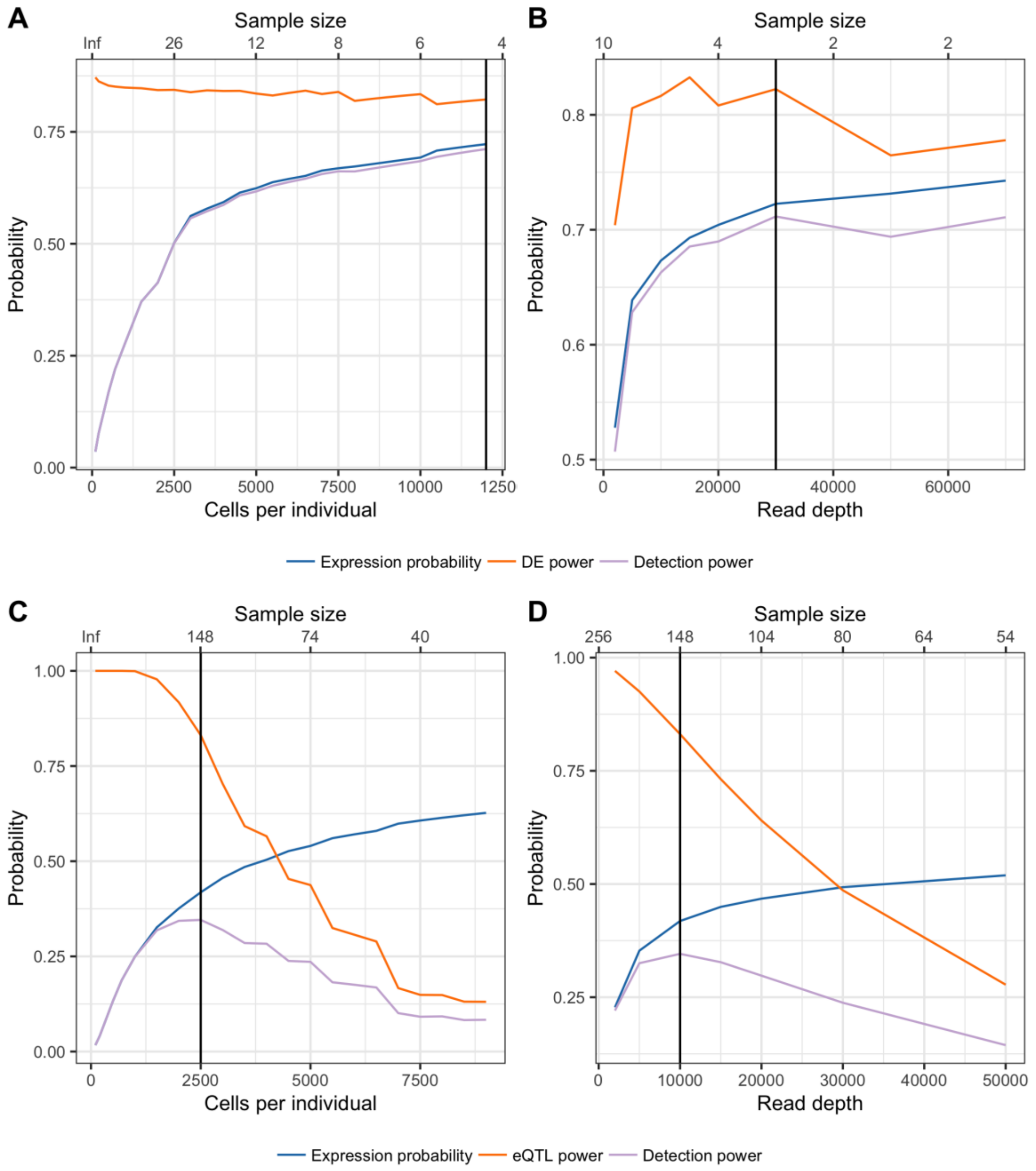


**Figure S11**: *Parameter optimization for constant budget*. Optimized parameters (sample size, read depth and cells per individual) to maximize the detection power for a DE study with a budget of 10,000€ (A,B) and an eQTL study with a budget of 30,000€ (D,E), both for a cell type with a frequency of 25%. Suplots A and C show the influence of the cells per individual given the optimized read depth, subplots B and D show the influence of the cells per individual given the optimized number of cells per individual. The third parameter, the sample size, is defined uniquely given the other two parameters due to the budget restriction. The optimal sample size values are shown in the upper x axes (no linear scale, but matching numbers to the parameter on the lower y axis under the given budget). The horizontal line in the subplots visualizes the optimal parameter combination. The same effect sizes and expression definition as in Figure 3 was taken.

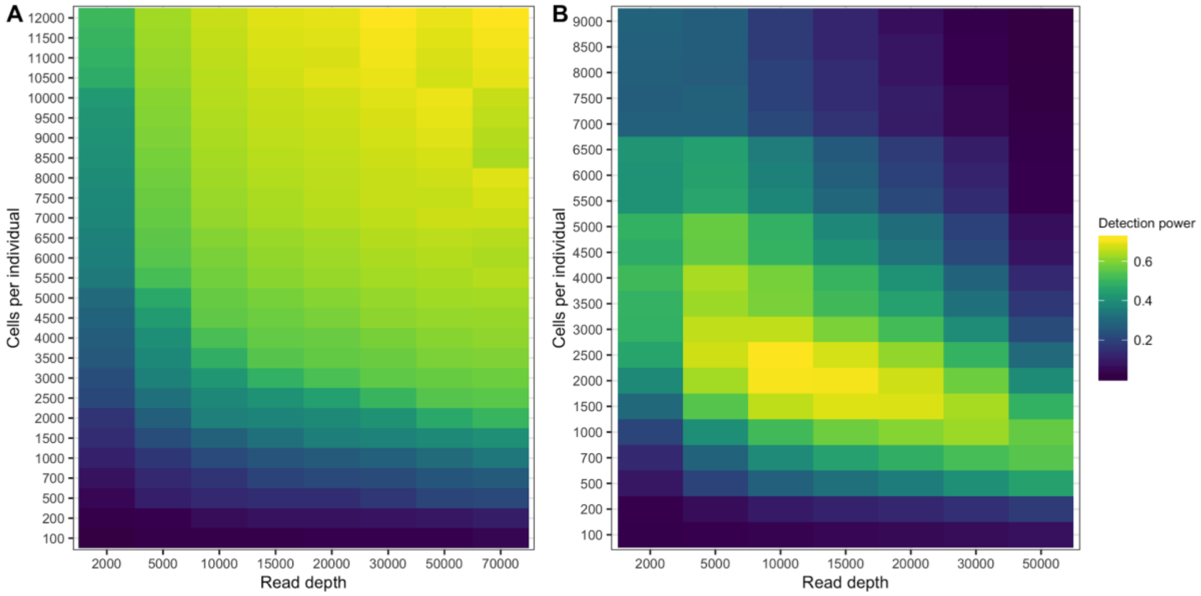


**Figure S12**: *Parameter optimization grid*. Maximizing detection power by selecting the best combination of cells per individual and read depth for A. a DE study with a budget of 10,000€ and B. a eQTL study with a budget of 30,000€, both for a cell type with a frequency of 25%. The sample size is defined uniquely given the other two parameters due to the budget restriction. The same effect sizes and expression definition as in Figure 3 were taken.

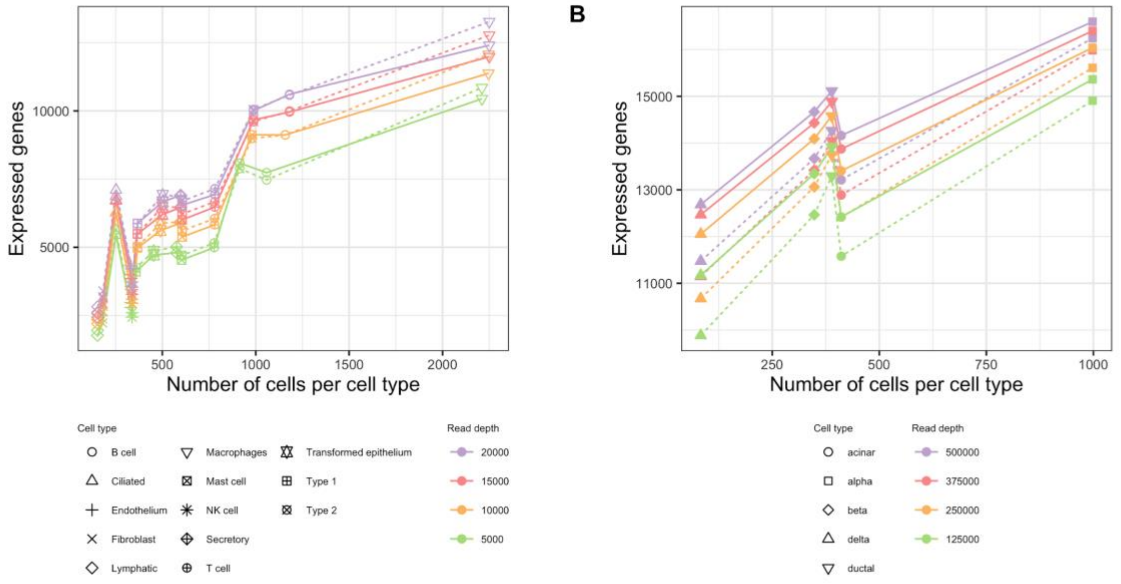


**Figure S13**: *Gene curve fits for different single cell technologies*. Plot A shows the gene curve fit for a lung data set measured with Drop-seq and plot B for a pancreas data set measured with Smart-seq2, both subsampled to different read depths. The solid lines represent the observed gene curves, the dashed lines the fitted curves. The point symbol visualizes the cell type. The definition for an expressed gene was parameterized in the following way: in Plot A, the gene needs to have UMI counts > 10 in all measured cells in total, in Plot B, read counts > 10 per kilobase transcript in all measured cells in total.

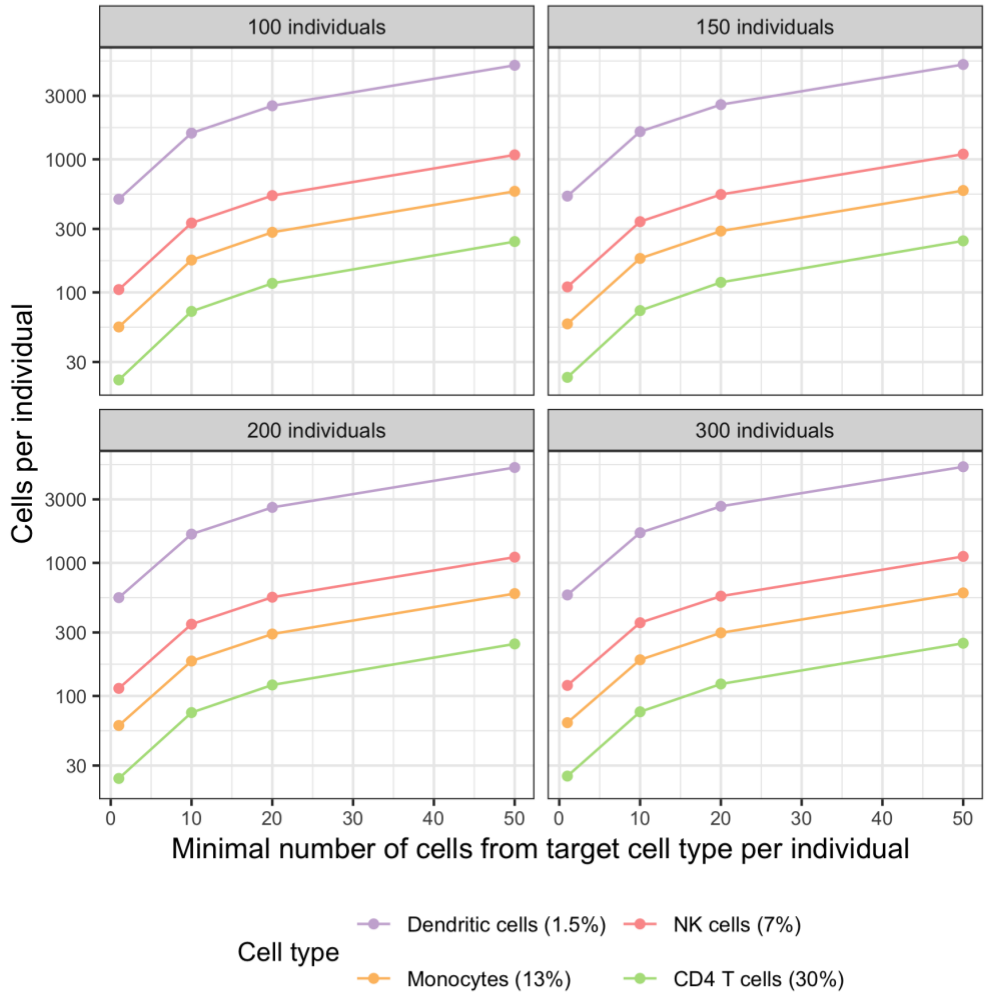


**Figure S14**: *Power to detect rare cell types*. The figure shows the required number of cells per individual (y-axis, log scale) to detect the minimal number of cells from a target cell type per individual (x-axis) with a probability of 95%. The probability depends on the total number of individuals and the frequency of the target cell type (purple, red, yellow, green). Note that the required number of cells per sample only counts “correctly measured” cells (no doublets etc), so the number is a lower bound for the required cells to be sequenced.

### Supplementary Tables

| Run | Donors | Target cell number | Target reads per cell | Estimated number of cells | Mean reads per cell |
|---|---|---|---|---|---|
| Run 1 | 1-14 | 8,000 | 50,000 | 7,491 | 40,650 |
| Run 2 | 1-7 | 8,000 | 50,000 | 5,989 | 127,685 |
| Run 3 | 8-14 | 8,000 | 50,000 | 8,144 | 13,949 |
| Run 4 | 1-14 | 8,000 | 50,000 | 7,429 | 35,417 |

| | | | | | |
|---|---|---|---|---|---|
| Run 5 | 1-14 | 8,000 | 50,000 | 7,765 | 21,057 |
| Run 6 | 1-14 | 25,000 | 50,000 | 20,126 | 51,792 |

**Table S1**: *PBMC data set*. Experimental parameters of the 6 PBMC runs. In Run 1, 4, 5 and 6 all 14 donors were measured, in Run 2 only donor 1-7 and in Run 3 only donor 8-14. Run 6 was overloaded with 25,000 cells. The estimated number of cells and mean reads per cell are taken from the cell ranger summary statistics.

| Cell type | Markers |
|---|---|
| CD4 T cells | IL7R, CD3D |
| CD14+ Monocytes | CD14, LYZ |
| B cells | MS4A1, CD79A |
| CD8 T cells | CD8A, CD8B, CD3D |
| NK cells | GNLY, NKG7 |
| FCGR3A+ Monocytes | FCGR3A, MS4A7 |
| Dendritic cells | FCER1A, CST3 |
| Megakaryocytes | PPBP |
| Plasma cells | CD79A |

**Table S2**: *Marker genes*. Marker genes used to assign the Louvain clusters to the cell types. Annotations taken from van der Wijst et al., 2018 [23] and the Scanpy PBMC tutorial [82].

| Technology | Library preparation costs per cell | Sequencing costs per 1 million reads |
|---|---|---|
| 10X Genomics | 0.05 € - 0.12€ | 3.42 € |
| Drop-Seq | 0.09 € | 3.42 € |
| Smart-Seq2 | 13.00 € | 3.42 € |

**Table S3**: *Experimental cost per technology*. Library preparation cost estimation (per cell) and sequencing cost estimation (per 1 million reads) for three of the most common single cell RNA-seq technologies in Euro (€). For 10X Genomics, the cost depends on the number of cells per lane, an overloading of each lane with 20,000 cells generates costs of 0.05€ per cell, a loading with 8,000 cells per lane costs of 0.12€ per cell.
